# CBP/p300 lysine acetyltransferases inhibit HIV-1 expression in latently infected T-cells

**DOI:** 10.1101/2024.07.05.602286

**Authors:** Riley M. Horvath, Ivan Sadowski

## Abstract

HIV-1 latency is regulated by chromatin modifying enzymes, and histone deacetylase inhibitors (HDACi) were previously found to reactivate provirus expression. We report that inhibitors of CBP/p300 acetyltransferases also cause reversal of latency in T-cells. CBP/p300 inhibitors synergize with mechanistically diverse latency reversing agents to cause HIV-1 reactivation. In contrast, inhibition of CBP/p300 impaired the latency reversal by the HDACi SAHA, indicating that CBP/p300 contribute to acetylation on the HIV-1 LTR associated with HDACi-mediated latency reversal. CBP/p300 inhibition caused loss of H3K27ac and H3K4me3 from the LTR, but did not affect association of the inhibitor protein BRD4. Furthermore, inhibition of the additional lysine acetyl transferases PCAF/GCN5 or KAT6A/KAT6B also caused reversal of latency, suggesting that protein acetylation has an inhibitory effect on HIV-1 expression. Collectively, these observations indicate that transcription from the HIV-1 LTR is controlled both positively and negatively by protein acetylation, likely including both histone and non-histone regulatory targets.

## Introduction

The advent of combination antiretroviral therapy (cART) has greatly improved the prognostic outcome of human immunodeficiency virus type 1 (HIV-1) infection by effectively managing the retrovirus as a chronic illness. However, cART does not represent a cure and must be administered lifelong as viral rebound promptly ensues following disruption of treatment in people living with HIV-1 (PLWH) (Chun et al. 2000) (SMART Study Group et al. 2006). The predominant barrier to a cure for HIV-1 is the extremely long-lived CD4^+^ T-cells that possess chromosomally integrated but transcriptionally silent latent provirus that develop over the course of infection. This latent population serves as a reservoir of viremia that escapes immune system surveillance while remaining capable of reactivation leading to recurrent seeding of the HIV-1 infection (Chun et al. 1998). HIV-1 infection persists in PLWH because of latently infected cells, and consequently much attention has been placed on manipulating proviral expression with the objective of eradicating these cells or blocking stochastic reactivation in order to eliminate the requirement of continuous cART administration. Two seemingly antithetical approaches have been proposed towards potential cures, designated the ‘shock and kill’ and the ‘block and lock’ strategies. The proposed ‘shock and kill’ tactic would involve application of latency reversing agents (LRAs) to induce proviral expression enabling removal of infected cells through a combination of immune system clearance and viral mediated cytotoxicity (Deeks, 2012) (Margolis et al. 2020). On the other hand, the ‘block and lock’ approach would involve intervention with latency promoting agents (LPAs) to enforce deep latency whereby spurious proviral reactivation is inhibited in the absence of ongoing treatment (Lewis et al. 2023) (Mediouni et al. 2022).

The ‘shock and kill’ strategy has been intensely investigated and has led to identification of diverse LRAs through both mechanism-based approaches and small molecule compound screens (Stoszko et al. 2019) (Ait-Ammar et al. 2020) (Hashemi & Sadowski, 2020). Multiple classes of LRAs have been described that include T-cell signaling agonists (Fromentin et al. 2019), kinase agonists (Díaz et al. 2015) (Jiang et al. 2015), epigenetic modifiers (Contreras et al. 2009) (Bernhard et al. 2011) (Marian et al. 2018) (Prins et al. 2023), transcriptional elongation enhancers (Soliman et al. 2023), bromodomain and extra-terminal domain (BET) inhibitors (Zhu et al. 2012) (Li et al. 2013) (Boehm et al. 2012), and TRIM24 bromodomain inhibitors (Horvath et al. 2023) (Horvath et al. 2023). Several barriers to the “shock and kill” strategy have become apparent, including the need to penetrate diverse tissue types that harbor HIV-1 infected cells (i.e. anatomical sanctuary sites), the potential inability to provoke a sufficiently large proviral reactivation response (Siliciano & Siliciano, 2021), and the inefficient elimination of infected cells by CD8^+^ killer T-cells (Gunst et al. 2020) (Powell et al. 2020). In general, a greater understanding of the mechanisms behind HIV-1 latency is required to produce a successful HIV-1 elimination strategy using this approach.

Given the obstacles to ‘shock and kill’, an alternative ‘block and lock’ strategy has garnered more recent attention, where the intention is to use LPAs to enforce production of deep latency where provirus expression remains suppressed in the absence of cART (Lewis et al. 2023) (Mediouni et al. 2022). One well characterized LPA is Didehydro-Cortistatin A (dCa), an analog of a natural steroidal alkaloid from the marine sponge *Corticium simplex* (Aoki et al. 2006). dCA was found to inhibit the viral transactivator Tat through direct interaction resulting in formation of a restrictive LTR epigenetic composition that limits RNAPII recruitment (Mousseau et al. 2012) (Mousseau et al. 2015) (Li et al. 2019). Importantly, HIV-1 suppression is maintained following the discontinuation of dCA treatment (Kessing et al. 2017) (Mediouni et al. 2019). Somewhat similar to dCA, the Nullbasic Tat mutant disrupts the Tat/P-TEFb positive feedback loop resulting in suppression of proviral transcription (Jin et al. 2016). More recently, inhibitors of the Mediator Complex kinases, CDK8 and CDK19, have been shown to repress HIV-1 reactivation from latency (Horvath et al. 2023). Furthermore, the CDK8/19 inhibitors Senexin A and BRD6989 suppress stochastic proviral expression, an effect that is maintained following discontinuation of CDK8/19 inhibition (Horvath et al. 2024). Although additional LPAs are continually being discovered, clinical trials have yet to be initiated for this potential therapeutic strategy.

The silencing of HIV-1 expression occurs *via* epigenetic processes (Nguyen & Karn, 2024). HIV-1 is a retrovirus that predominately integrates into actively transcribed chromatin and becomes subject to regulation by histone modifying enzymes that promote an active or restrictive epigenetic environment on the LTR promoter (Battivelli et al. 2018). Generally, histone demethylases (HDMs) compete with the activity of histone methyltransferases (HMTs) for methylation of histone N-terminal tail lysine methylation, while histone acetyltransferases (HATs) antagonize the activity of histone deacetylases (HDACs). Histone methylation is known to promote or restrict transcription dependent on the target lysine residue. For instance, H3K27me is associated with transcriptional repression while H3K4me correlates with activation. Methylation of H3K9, H3K27, and H4K20 have been shown to be associated with proviral latency where inhibition of the catalyzing HMTs cause latency reversal (Imai et al. 2010) (Friedman et al. 2011) (Bernhard et al. 2011) (Boehm et al. 2017) (Nguyen et al. 2017) (Ye et al. 2022) (Lu et al. 2022).

Additionally, transcription from the provirus is repressed by multiple HDACs, which are recruited by factors such as YY1, NFκB p50, and CBF-1 (Verdikt et al. 2021) (Peterson et al. 2023), which remove acetylation from LTR-associated histones. Notably, histone deacetylase inhibitors (HDACi) are a predominant LRA class that have already been evaluated in clinical trials (Kumar et al. 2015) (Wightman et al. 2012), but to date have not reduced the size of the latent reservoir (Siliciano et al. 2007) (Archin et al. 2008) (Sagot-Lerolle et al. 2008) (Archin et al. 2010) (Rasmussen et al. 2014) (McMahon et al. 2021) (Gay et al. 2022). Opposing HDAC activity are the histone/lysine acetyl transferases (HATs/KATs), including CBP/p300, which are recruited to the HIV-1 LTR upon transcriptional induction, and whose presence correlates with enrichment of histone acetylation (Lusic et al. 2003). In particular, NFAT and Tat have been shown to facilitate the recruitment of CBP and its paralog p300 to the transcriptionally active LTR (Benkirane et al. 1998) (García-Rodríguez & Rao, 1998) (Lusic et al. 2003). However, evidence of the direct role of CBP/p300 for the regulation of HIV-1 expression by specific acetylation of target protein(s) is lacking.

Given the association of histone acetylation and activation of HIV-1 provirus, we examined whether inhibition of CBP/p300 acetyltransferase activity would inhibit provirus reactivation, using the recently developed small molecule inhibitors A-485 (Lasko et al. 2017) and iP300w (Bosnakovski et al. 2019). To our surprise, inhibition of CBP/p300 activity with these compounds induced HIV-1 transcription in both cell line models of latency and primary CD4^+^ PBMCs *ex vivo*. When used in combination with either the MAPK/PKC agonist PMA, the NFκB activator PEP005, the BET bromodomain inhibitor JQ1, or the TRIM24 bromodomain inhibitor IACS-9571, CBP/p300 inhibition displayed synergistic and robust induction of proviral expression. Consistent with the role of CBP/p300 for acetylation dependent gene activation, both A-485 and iP300w antagonized latency reversal in response to the HDACi SAHA. Furthermore, PROTAC mediated degradation of CBP/p300 using dCBP-1 mirrored the effects of the chemical inhibitors, confirming the latency promoting function of these factors (Vannam et al. 2021). In addition, although CBP/p300 inhibition induced HIV-1 expression, we observed the loss of H3K27ac and H3K4me3 epigenetic marks at the proviral LTR, which are generally associated with activation. This work reveals that acetylation of protein(s) by CBP/p300 causes an inhibitory effect HIV-1 LTR, opposite that of known effects of histone acetylation for regulation of transcription. This unique effect requires consideration for development of therapeutic strategies to modulate HIV-1 expression, particularly involving HDAC inhibitors.

## Results

### CBP/p300 acetyltransferase inhibitors induce HIV-1 transcription

Given the presumption that CBP/p300 acetyltransferases are associated with reactivation of proviral latency (Mbonye & Karn, 2017) (Ne et al. 2018) (Moranguinho & Valente, 2020) (Nguyen & Karn, 2024), we sought to characterize the effect of CBP/p300 chemical inhibitors on HIV-1 transcription. To this end, we treated the JLat10.6 T-cell model of HIV-1 latency with A-485 (Lasko et al. 2017) and iP300w (Bosnakovski et al. 2019), two recently developed small molecules that are highly specific inhibitors of CBP/p300 acetyltransferase activity (Fig. 1*A*). The JLat10.6 cell line possesses a chromosomally integrated provirus where GFP is inserted in place of *nef* and serves as a readout for HIV-1 5’ LTR expression, while a frameshift in *env* prevents additional rounds of replication (Jordan et al. 2003). To our surprise, we observed that the inhibitors caused a dose-dependent increase in HIV-1 expression as measured by 5’ LTR driven GFP expression (Fig. 1*B*, 1*C*). Treatment of cells with A-485 and iP300w for 24 hrs produced a modest 2.7- and 3.1-fold induction in HIV-1 expression respectively (Fig. S1, 24 hrs), but expression increased considerably after two days treatment to 6.5x for A-485 and 8.9x for iP300w (Fig. S1, 48 hrs). Importantly, the inhibitors did not generate autofluorescence (Fig. S2) and cell viability was unaffected at the concentrations applied (Fig. 1*D*).

**Figure 1.**
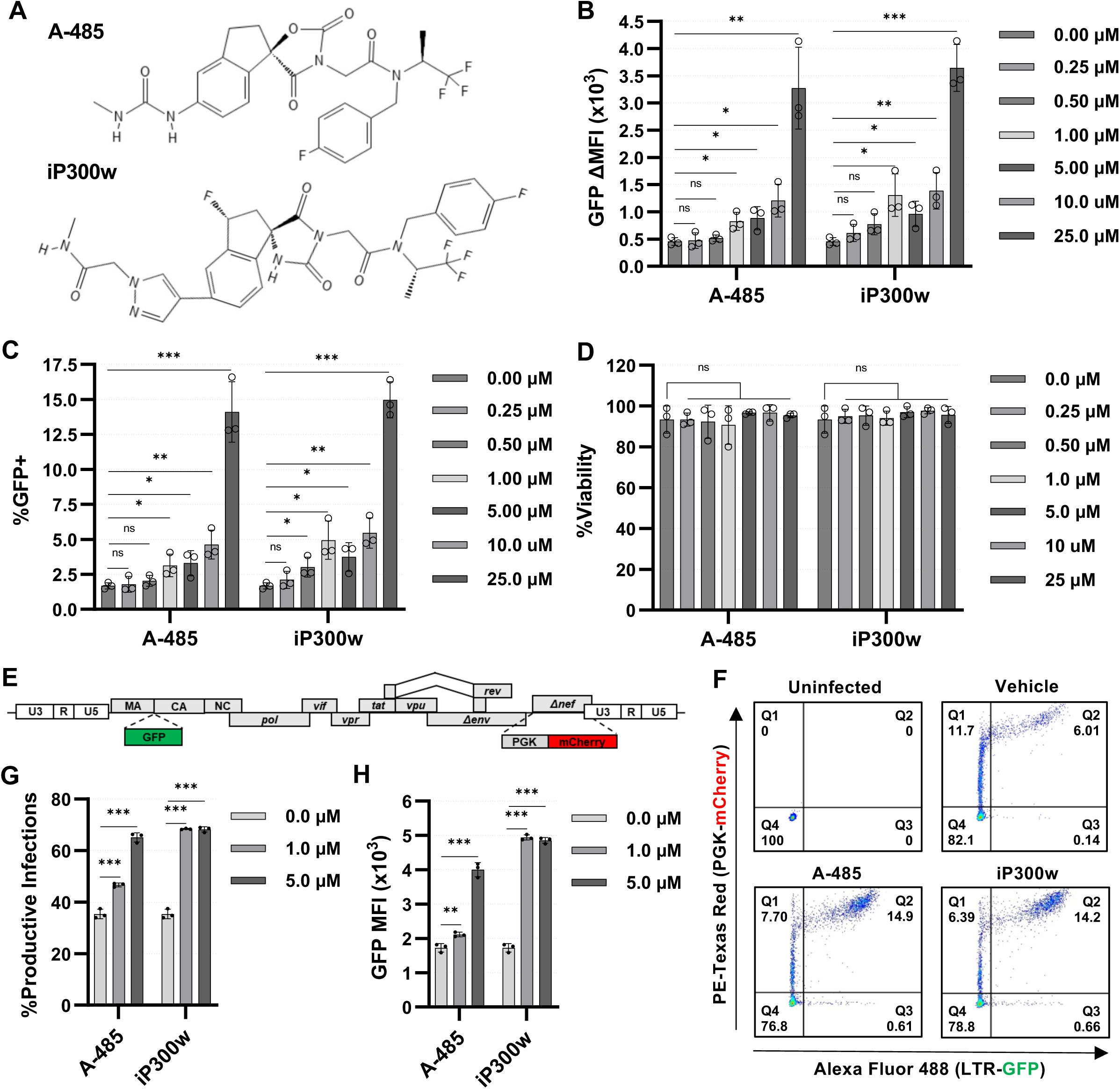
Inhibitors of CBP/p300 acetyltransferase activity promote HIV-1 transcription. **A:** Molecular structure of the highly selective CBP/p300 acetyltransferase inhibitors A-485 and iP300w. **B**, **C:** JLat10.6 cells were incubated with the indicated concentration of CBP/p300 inhibitor for 24 hrs. Subsequently, HIV-1 expression was assessed by flow cytometry and is reported as the GFP delta (Δ) Mean Fluorescence Intensity (MFI) (B) and the proportion of GFP expressing cells (C) (*n* = 3, mean ± SD, unpaired *t*-test). **D:** Cellular viability was determined for JLat10.6 cells treated as in (B) and (C) (*n* = 3, mean ± SD, unpaired *t*-test). **E:** Schematic representation of the Red-Green-HIV-1 full-length dual reporter virus (Dahabieh et al, 2013). GFP expression is directed by the 5’ LTR and is a measure of proviral transcription, while an internal constitutive PGK promoter drives mCherry expression allowing for the determination of infection independent of LTR activity. A frameshift mutation in *env* renders the virus replication incompetent. **F:** Representative flow cytometry scatter plots of RGH infected Jurkat E6-1 T-cells that 1-day post-infection, were treated with a vehicle control (DMSO), 5 μM A-485, or 5 μM iP300w for 48 hrs. Latently infected cells are in Q1 (mCherry+), productive HIV-1 infected T-cells are in Q2 (GFP+/mCherry+), noise generated by viral recombination is in Q3 (GFP+), and uninfected cells are in Q4. **G, H:** Jurkat E6-1 T-cells were transduced with RGH at a multiplicity of infection that caused ∼20% of cells to be infected. 24 hrs post-infection, cells were incubated with a vehicle control (DMSO), or the indicated concentration of A-485 or iP300w. Following 48 hrs, HIV-1 expression was determined by flow cytometry and is reported as the percent of productively infected cells (Q2/(Q1+Q2)x100) (G) and the GFP Mean Fluorescence Intensity (MFI) of the mCherry+ population (Q1 and Q2) (H) (*n* = 3, mean ± SD, unpaired *t*-test).

Since the establishment of latency can occur early upon infection (Dahabieh et al. 2013) (Dahabieh et al. 2014) (Battivelli et al. 2018), we examined the effect of CBP/p300 inhibition following infection of T-cells with the Red-Green-HIV-1 (RGH) dual fluorescence reporter virus (Dahabieh et al. 2013). RGH is a full-length HIV-1 reporter where GFP is expressed from the 5’ LTR, and a constitutive promoter expressing mCherry is inserted in place of *nef* to allow differentiation between latent and productively infected cells (mCherry+/GFP-versus mCherry+/GFP+). Additionally, a frameshift mutation in *env* renders this virus replication incompetent, preventing multiple rounds of infection (Fig. 1*E*). Following infection of Jurkat CD4^+^ T-cells with RGH, treatment with CBP/p300 inhibitors caused an increase in the proportion of productive infections (Fig. 1*F*, 1*G*) and resulted in a significant increase in expression from the 5’ LTR in infected cells as measured by GFP intensity (Fig. 1*H*). Collectively, these results suggest that CBP/p300 acetyltransferase activity causes repression of HIV-1 expression and contributes to the development of latency upon infection.

### CBP/p300 inhibitors synergize with mechanistically distinct LRAs for HIV-1 reactivation

As CBP/p300 inhibition was found to reverse HIV-1 latency, we examined the effect of A-485 and iP300w in combination with various mechanistically distinct LRAs. Treatment of JLat10.6 T-cells with PMA in combination with a range of CBP/p300 inhibitor concentrations revealed robust and synergistic activation of HIV-1 expression as determined by Bliss Independence Modeling (Fig. 2*A*, 2*B*). Of note, both inhibitors generated a dose-dependent increase in HIV-1 expression that was greater than treatment with PMA alone (Fig. 2*A*), although lower concentrations of iP300w than A-485 were required to produce synergistic activation (Fig. 2*B*). Additionally, no effect on cellular viability was observed for any of the treatments (Fig. S3*A*).

**Figure 2.**
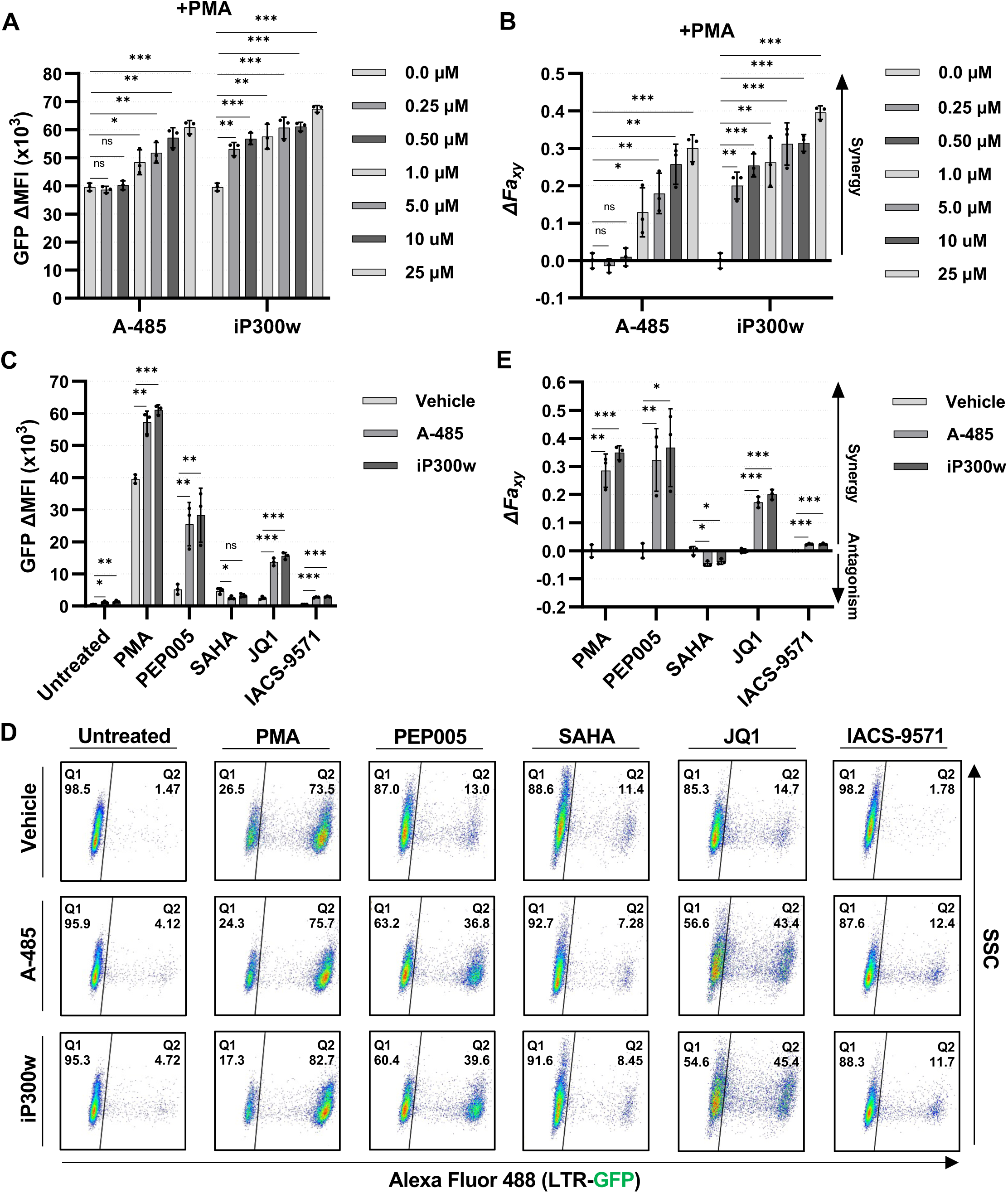
Chemical interaction between CBP/p300 inhibitors and various LRAs. **A:** JLat10.6 cells were pre-treated with the indicated concentration of A-485 or iP300w for 1 hr, after which 4 nM PMA was added. Following 24 hrs, HIV-1 expression was determined by flow cytometry and is indicated as the GFP delta (Δ) Mean Fluorescence Intensity (MFI) (*n* = 3, mean ± SD, unpaired *t*-test). **B:** Drug interaction between 4 nM PMA and the indicated concentration of either A-485 or iP300w was determined using Bliss Independence Modeling. Presented is the difference between the predicted and the observed fractional HIV-1 transcription response to the given drug combination whereby values greater than 0 indicate a synergistic interaction. See Materials and Methods for more details (*n* = 3, mean ± SD, unpaired *t*-test). **C:** JLat10.6 cells were treated with a vehicle control (DMSO), 10 μM A-485, or 10 μM iP300w for 1 hr prior to the addition of the indicated latency reversing agent. Following 24 hrs, HIV-1 expression was assessed by flow cytometry and is reported as the GFP delta (Δ) Mean Fluorescence Intensity (MFI). The concentration of LRAs used were 4 nM PMA, 4 nM PEP005, 1 μM SAHA, 10 μM JQ1, 15 μM IACS-9571 (*n* = 3, mean ± SD, unpaired *t*-test). **D:** Representative flow cytometry scatter plots for JLat10.6 cells treated as indicated in (C). Q1 contains latent (GFP-) cells while cells possessing transcriptionally active provirus are shown in Q2 (GFP+). **E:** Bliss Independence Modeling was applied to determine combinatorial drug interaction as in (B). Values greater than 0 indicate synergy while values less than 0 signify an antagonistic relationship (*n* = 3, mean ± SD, unpaired *t*-test).

Next, we compared the effect of CBP/p300 inhibitors in combination with the additional previously characterized LRAs PMA, PEP005, SAHA, JQ1, and IACS-9571. The phorbol ester PMA is a Ras-MAPK/PKC agonist that functions through activation of various factors including NFκB, AP1, and GABP/ETS; PEP005 causes PKC dependent NFκB activation; SAHA is a pan-HDACi; JQ1 is a BET protein (BRD4) bromodomain inhibitor; and IACS-9571 is a TRIM24 bromodomain inhibitor that stimulates HIV-1 transcriptional elongation (Horvath et al. 2023) (Horvath et al. 2023). We observed significant reversal of proviral latency for each combination of CBP/p300 inhibitor and LRA as compared to individual treatment, with the exception of SAHA where the combination with CBP/p300 inhibitor slightly inhibited transcriptional induction (Fig. 2*C*, 2*D*). Bliss Independence Modeling confirmed that CBP/p300 inhibitors cause a synergistic effect with PMA, PEP005, JQ1, and IACS-9571, but are antagonistic towards SAHA (Fig. 2*E*). Importantly, cellular toxicity was not observed for any combination of drugs examined (Fig. S3*B*). Because SAHA is an HDACi that reverses latency, presumably by enhancing histone acetylation, the finding that CBP/p300 inhibitors severely reduce SAHA mediated HIV-1 induction indicates that CBP/p300 must contribute to accumulation of protein acetylation associated with activation of transcription from the HIV-1 LTR. However, we note that multiple HATs must be required for the latency reversing effect of SAHA because CBP/p300 inhibitors only partially prevent reactivation. More importantly, because inhibition of CBP/p300 reverses proviral latency (Fig. 1) and synergizes with other LRAs for reactivation of HIV-1 provirus, additional uncharacterized protein acetylation must cause repression of HIV-1 transcription.

### PROTAC mediated CBP/p300 degradation reactivates latent HIV-1

Given the surprising finding that inhibitors of CBP/p300 acetyltransferase activity cause reactivation of HIV-1 expression, we sought to confirm this effect using dCBP-1, a proteolysis targeting chimera (PROTAC) consisting of a CBP/p300 bromodomain binding motif and the cereblon E3 ubiquitin ligase targeting moiety that causes the proteasomal degradation of CBP and p300 (Vannam et al. 2021). We first confirmed a range of dCBP-1 concentrations that induces the near total degradation of CBP and p300 proteins within 24 hours (Fig. 3*A*). Treatment of JLat10.6 T-cells with dCBP-1 caused dose-dependent induction of HIV-1 expression (Fig. 3*B*), the magnitude of which mirrored CBP/p300 acetyltransferase inhibition (Fig. 1*B*). Akin to the acetyltransferase inhibitors (Fig. S1), treatment with dCBP-1 over a week resulted in the gradual increase in HIV-1 expression, peaking at 31-fold greater than untreated cells (Fig. 3*C*). Next, we applied the CBP/p300 PROTAC in combination with LRAs to examine synergistic effects for reactivation of HIV-1 provirus. Similar to the above results with the CBP/p300 inhibitors, dCBP-1 in combination with PMA, PEP005, JQ1, or IACS-9571 generated greater HIV-1 expression than individual treatment, while causing a slight decrease in SAHA mediated latency reversal (Fig. 3*D*, 3*E*). As with A-485/iP300w treatment, we observed synergistic interactions between dCBP-1 and PMA, PEP005, JQ1, and IACS-9571, and an antagonistic relationship with SAHA (Fig. 3*F*). Treatment with dCBP-1 alone or in combination with the LRAs had no effect on cellular viability (Fig. S4). Taken together, these results confirm a repressive function of CBP/p300 on HIV-1 expression.

**Figure 3.**
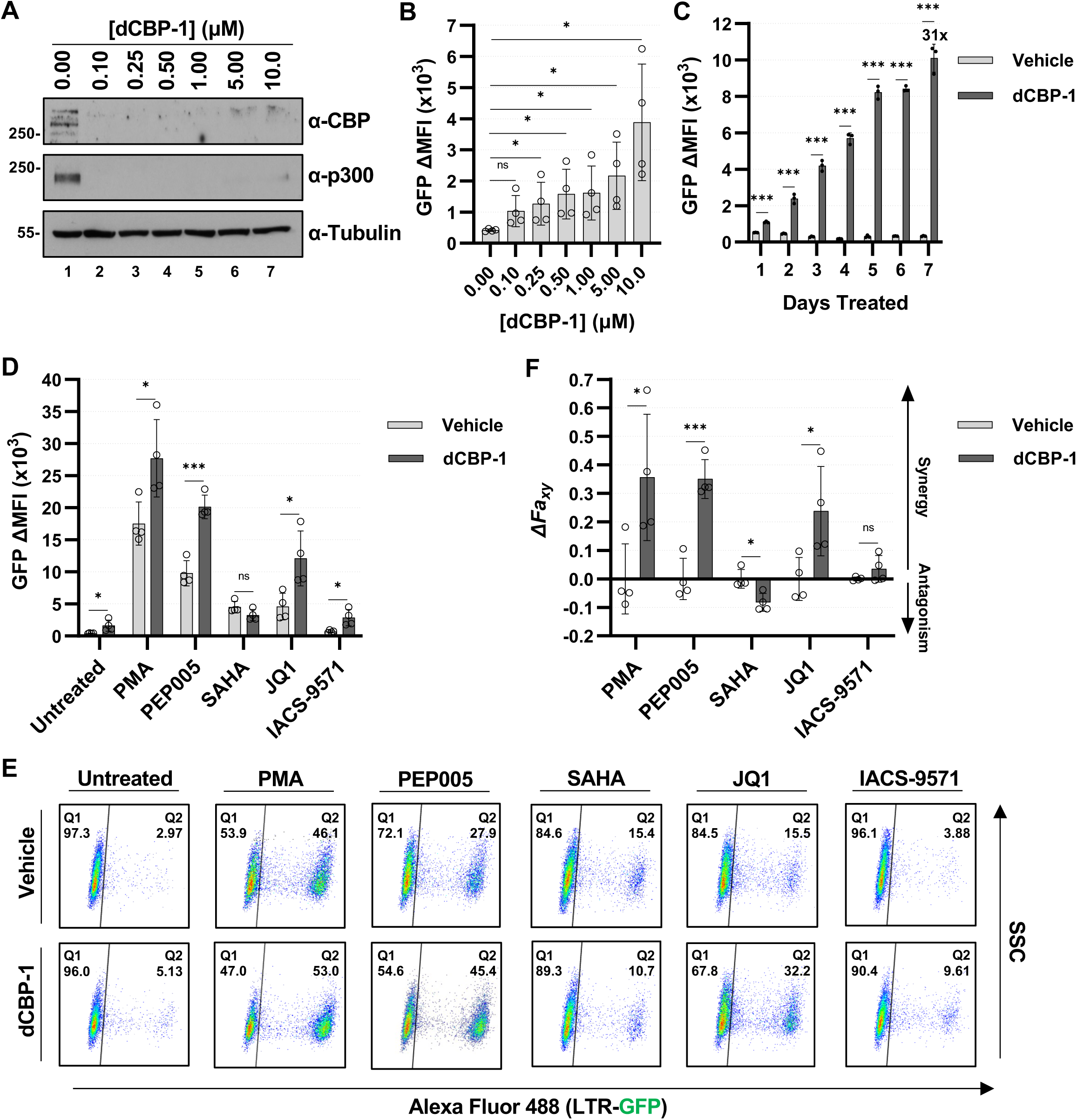
PROTAC mediated degradation of CBP/p300 reactivates latent HIV-1. **A:** JLat10.6 cells were incubated with the indicated concentration of dCBP-1. Following 24 hrs, whole cell lysates were extracted and subjected to immunoblotting with the indicated antibody. **B:** JLat10.6 cells were incubated for 24 hrs with the indicated concentration of dCBP-1. After 24 hrs, HIV-1 expression was assessed by flow cytometry and is depicted as the change (Δ) in GFP Mean Fluorescence Intensity (MFI) (*n* = 4, mean ± SD, unpaired *t*-test). **C:** JLat10.6 cells were incubated with a vehicle control or 1 μM dCBP-1. Following the indicated amount of time, HIV-1 expression was assessed by flow cytometry and is reported as the GFP delta (Δ) Mean Fluorescence Intensity (MFI) (*n* = 3, mean ± SD, unpaired *t*-test). **D:** JLat10.6 cells were incubated in the presence of 1 μM dCBP-1 for 3 hrs. Subsequently, the indicated LRA was added, and HIV-1 expression was examined 24 hrs later by flow cytometry and is depicted as the delta (Δ) GFP Mean Fluorescence Intensity (MFI). The concentrations of LRAs used were 4 nM PMA, 4 nM PEP005, 1 μM SAHA, 10 μM JQ1, 15 μM IACS-9571 (*n* = 4, mean ± SD, unpaired *t*-test). **E:** Representative flow cytometry scatter plots for JLat10.6 cells treated as indicated in (D). Q1 contains latent (GFP-) cells while the transcriptionally active population is present in Q2 (GFP+). **F:** Bliss Independence Modeling was applied to determine combinatorial drug interactions between dCBP-1 and the indicated LRA. Values greater than 0 indicate synergy while values less than 0 signify an antagonistic relationship (*n* = 4, mean ± SD, unpaired *t*-test).

### CBP/p300 inhibition induces HIV-1 expression ex vivo

We also assessed the effect of CBP/p300 inhibition on HIV-1 expression *ex vivo*. For this, we infected primary CD4^+^ T-cells isolated from a healthy individual with the RGH dual reporter virus. 3 days post-infection, the cells were treated with either A-485 or iP300w for 24 hrs (Fig. 4*A*). Following treatment with CBP/p300 inhibitor, we observed a marginal increase in the proportion of productive infections (Fig. 4*B*, 4*C*) and ∼2-fold increase in expression from the 5’ LTR as measured by GFP intensity (Fig. 4*D*). Treatment with the inhibitors did not have a measurable impact on cellular viability (Fig. 4*E*).

**Figure 4.**
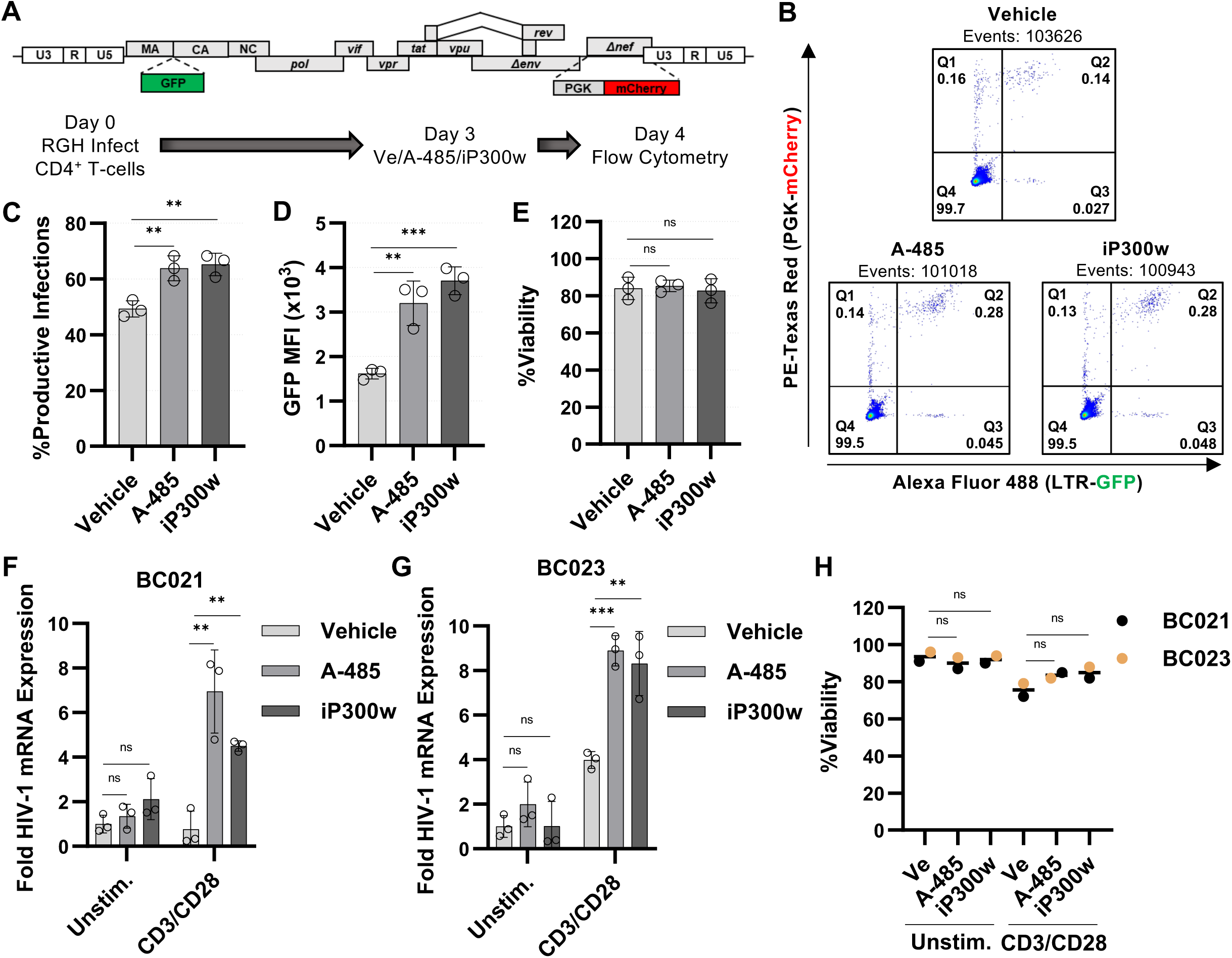
CBP/p300 inhibition induces HIV-1 expression in CD4^+^ cells *ex vivo*. **A:** Experimental design for RGH infection of primary CD4^+^ T-cells. CD4^+^ T-cells isolated from an uninfected participant were infected with the RGH dual reporter virus at a low MOI. 3 days, post-infection, cells were incubated with a vehicle control (DMSO), 10 μM A-485, or 10 μM iP300w for 24 hrs after which proviral expression was assessed by flow cytometry. **B:** Representative flow cytometry scatter plots of primary CD4^+^ T-cells treated as in (A). Latent infections are in Q1 (mCherry+), productively infected T-cells are in Q2 (GFP+/mCherry+), noise generated by viral recombination is in Q3 (GFP+), and uninfected cells are in Q4. **C, D:** Following treatment of primary CD4^+^ T-cells as in (A), proviral transcription was determined by flow cytometry and is expressed as the percentage of productive infections (C) and the GFP Mean Fluorescence Intensity (MFI) of infected cells (D) (*n* = 3, mean ± SD, one-way ANOVA with Dunnett’s multiple comparisons test). **E:** Viability was determined for participant derived CD4^+^ T-cells treated as in (A) (*n* = 3, mean ± SD, one-way ANOVA with Dunnett’s multiple comparisons test). **F, G:** Primary CD4^+^ PBMCs isolated from people living with HIV-1 who are receiving ART were treated with a vehicle (DMSO), 10 μM A-485, or 10 μM iP300w and were either left unstimulated or were treated with anti-CD3/anti-CD28. Following 24 hrs, intracellular RNA was extracted, and RT-PCR was preformed using oligos specific for multiply spliced Tat-Rev HIV-1 mRNA transcripts. HIV-1 mRNA expression is normalized to *GAPDH* (*n* = 3, mean ± SD, unpaired *t*-test). **H:** Primary CD4^+^ PBMCs treated as in (F) and (G) were assessed for cellular viability (*n* = 2, mean, unpaired *t*-test).

To further evaluate the role of CBP/p300 for HIV-1 latency, we tested the propensity of CBP/p300 inhibition to reactivate HIV-1 in CD4^+^ peripheral blood mononuclear cells (PBMCs) obtained from two aviremic participants who have been receiving cART. Treatment of cells from either participant with the CBP/p300 inhibitors on their own for 24 hrs did not produce a significant increase in HIV-1 mRNA (Fig. 4*F*, 4*G*, Unstim.). Latent proviruses can be induced by stimuli such as cytokine or antigenic stimulation that causes cellular activation through the T-cell receptor (TCR). In order to illicit such a response, we stimulated the participant CD4^+^ cells with CD3/CD28 antibodies which cross link and activate the T-cell receptor, alone or in combination with the CBP/p300 inhibitors. In this analysis we observed that stimulation by CD3/CD28 crosslinking produced a detectable increase in HIV-1 mRNA for participant BC023 but not BC021 (Fig. 4*F*, 4*G*, CD3/CD28, Vehicle). However, stimulation with CD3/CD28 antibodies in combination with either A-485 or iP300w produced a significantly greater increase in HIV-1 mRNA for both participants (Fig. 4*F*, 4*G*). In addition, we did not observe cellular toxicity in these cells upon addition of CBP/p300 inhibitors (Fig. 4*H*). For people living with HIV-1 (PLWH) who are receiving cART, studies indicate that roughly 1 per 10^6^ CD4^+^ T-cells possess a replication competent proviral genome (Shan & Siliciano, 2013). That being the case, it is possible that longer treatment intervals with CBP/p300 inhibitors alone would reveal a detectable change in HIV-1 mRNA. Nevertheless, these results indicate that CBP/p300 activity contributes to repression of HIV-1 provirus in PLWH.

### CBP/p300 inhibition does not cause global T-cell activation

Stimulation of the TCR through engagement with antigen presenting cells stimulates T-cell activation through the activity of TCR-associated protein tyrosine kinases which trigger the MAPK cascade, PKC, and calcineurin pathways (Sadowski & Mitchell, 2005). These pathways activate downstream transcription factors including NFκB, NFAT, AP1, and GABP/ Ets, which coordinate the inflammatory immune response by inducing expression of their respective target genes (Rothenberg, 2014). Intriguingly, the HIV-1 LTR has evolved to possess binding sites for many TCR induced transcription factors, including those mentioned above, thus coupling proviral transcription to T-cell activation. Consequently, a major challenge to the ‘shock and kill’ therapeutic strategy is to force proviral transcription without promoting global T-cell activation which can cause cytokine release syndrome, a life-threatening condition (Shimabukuro-Vornhagen et al. 2018).

In order to determine the effect of CBP/p300 inhibition on T-cell activation, we assessed available RNA-seq data to determine genes that are differentially regulated upon PMA/ionomycin stimulation (Horvath et al. 2023). This analysis identified 1481 downregulated genes and 2442 upregulated genes in activated CD4^+^ T-cells relative to unstimulated (Fig. S5). Notably, neither CBP (*CREBBP*) nor p300 (*EP300*) expression was found to be altered (Fig. S5). From this analysis, we selected six genes encoding cytokines and surface receptors including *IL2* and *CD69* which are known to be upregulated upon activation (Haas et al. 2011) (Fig. 5*A*). Using RT-PCR, we confirmed that PMA/ionomycin mediated T-cell activation induced expression of all six of the selected genes (Fig. 5*B*-*G*, PMA/Ion). Interestingly, we observed that CBP/p300 inhibition caused a range of effects on these genes. CBP/p300 inhibition was found to repress *IL2* and *CD69* expression, two genes that were strongly induced by T-cell activation (Fig. 5*B*, 5*C*). However, both A-485 and iP300w caused activation of *IL8* and *CSF2* expression, though these genes are only moderately upregulated (Fig. 5*D*, 5*E*). Finally, we found that the CBP/p300 inhibitors had minimal effect on the expression of *CCL22* and *XCL1* (Fig. 5*F*, 5*G*). These results indicate that CBP/p300 inhibition does not initiate global T-cell activation and likely does not promote HIV-1 latency reversal by co-opting a master immune response factor.

**Figure 5.**
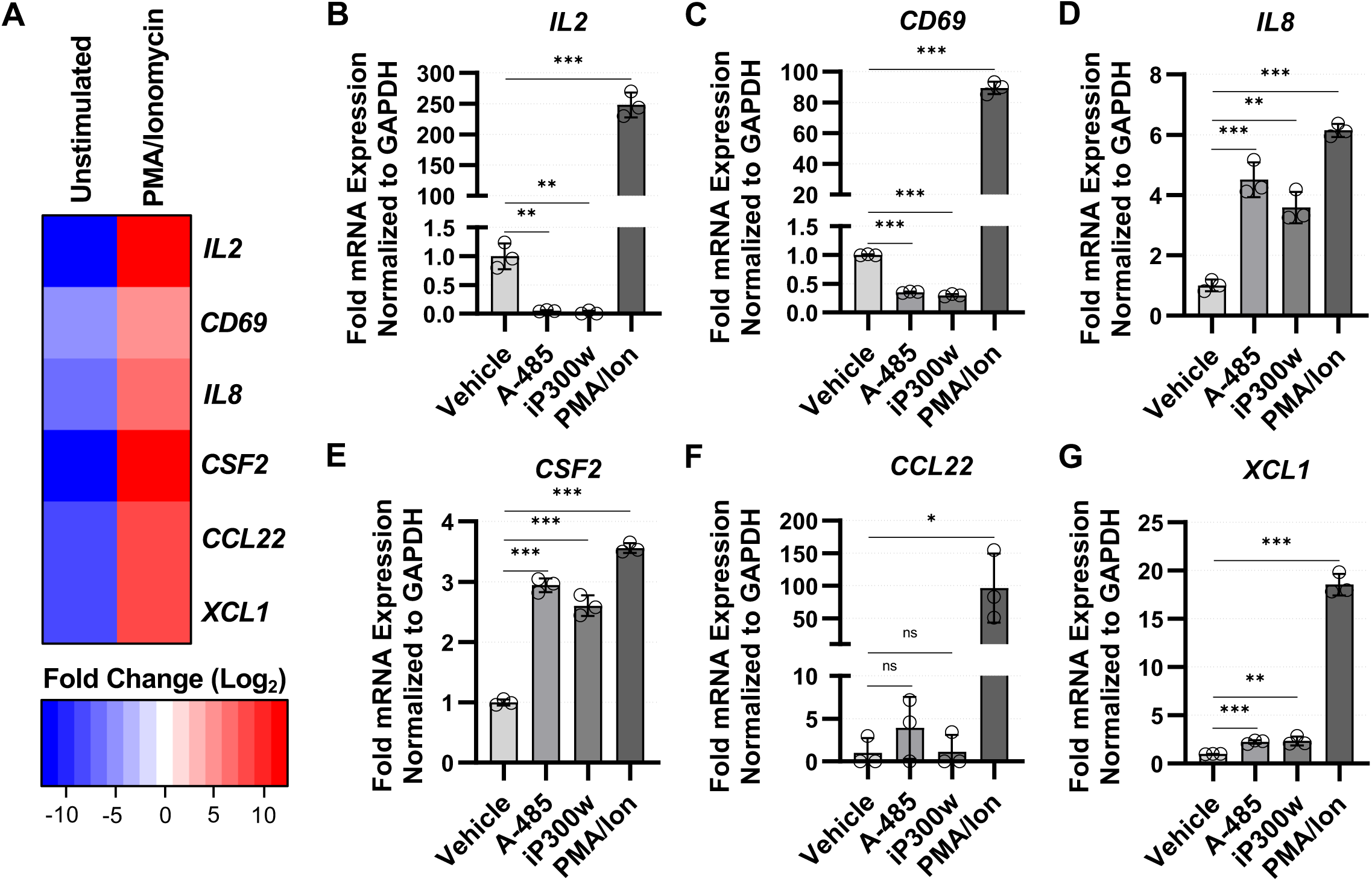
Effect of CBP/p300 inhibition on T-cell activation. **A:** Heatmap depiction of several upregulated genes following PMA/ionomycin treatment of Jurkat T-cells as identified by RNA-seq analysis (Horvath et al. 2023). **B – G:** Jurkat T-cells were incubated 24 hrs in the presence of DMSO (Vehicle), 10 μM A-485, 10 μM iP300w, or 4 nM PMA/ 1 μM ionomycin. Subsequently, intracellular RNA was extracted and analyzed by RT-PCR using oligos specific for the indicated mRNA transcript. RNA expression is normalized to *GAPDH* transcript (*n* = 3, mean ± SD, unpaired *t*-test).

### Dependence of Tat for regulation of HIV-1 by CBP/p300

The viral transactivator of transcription Tat is expressed from multiply spliced sub-genomic HIV-1 mRNA and produces a strong positive feedback loop by recruiting P-TEFb, a heterodimer of CDK9 and Cyclin T1, to the 5’ LTR through interaction with the nascent TAR RNA, which facilitates robust escape of promoter proximal paused RNA polymerase II (RNAPII) (Karn & Stoltzfus, 2012). Furthermore, CBP/p300 recruitment to the LTR has been shown to be facilitated by Tat, and CBP/p300 acetylation of Tat is thought to increase its transcriptional activator function (Benkirane et al. 1998) (García-Rodríguez & Rao, 1998) (Marzio et al. 1998) (Ott et al. 1999) (Lusic et al. 2003). Given the role of Tat for HIV-1 expression and the observed interplay with CBP/p300, we examined the dependency of Tat for HIV-1 reactivation in response to CBP/p300 inhibition using the JLatA72 cell line. This Jurkat derived CD4^+^ T-cell line harbors a chromosomally integrated HIV-1 LTR that controls GFP expression independently from Tat (Jordan et al. 2001). Treatment of JLatA72 cells with either A-485 or iP300w caused induction of GFP expressed from the LTR, indicating that Tat is not required for stimulation of HIV-1 expression by the CBP/p300 inhibitors (Fig. 6*A*, 6*B*, Untreated). Additionally, combined treatment of the CBP/p300 inhibitors with PMA, PEP005, JQ1, or IACS-9571 generated much higher levels of HIV-1 transcription than any treatment alone (Fig. 6*A*, 6*B*), but CBP/p300 inhibition somewhat restricted latency reversal in response to the HDACi SAHA in the JLatA72 line (Fig. 6*A*, 6*B*, SAHA). Bliss Independence Modeling demonstrated similar relationships as described above, although the synergistic interaction between the CBP/p300 inhibitors and JQ1 was less in JLatA72 cells than that for the JLat10.6 cell line (Fig 2*E*, Fig. 6*C*). Collectively, these results indicate that CBP/p300 inhibits HIV-1 expression independently of Tat.

**Figure 6.**
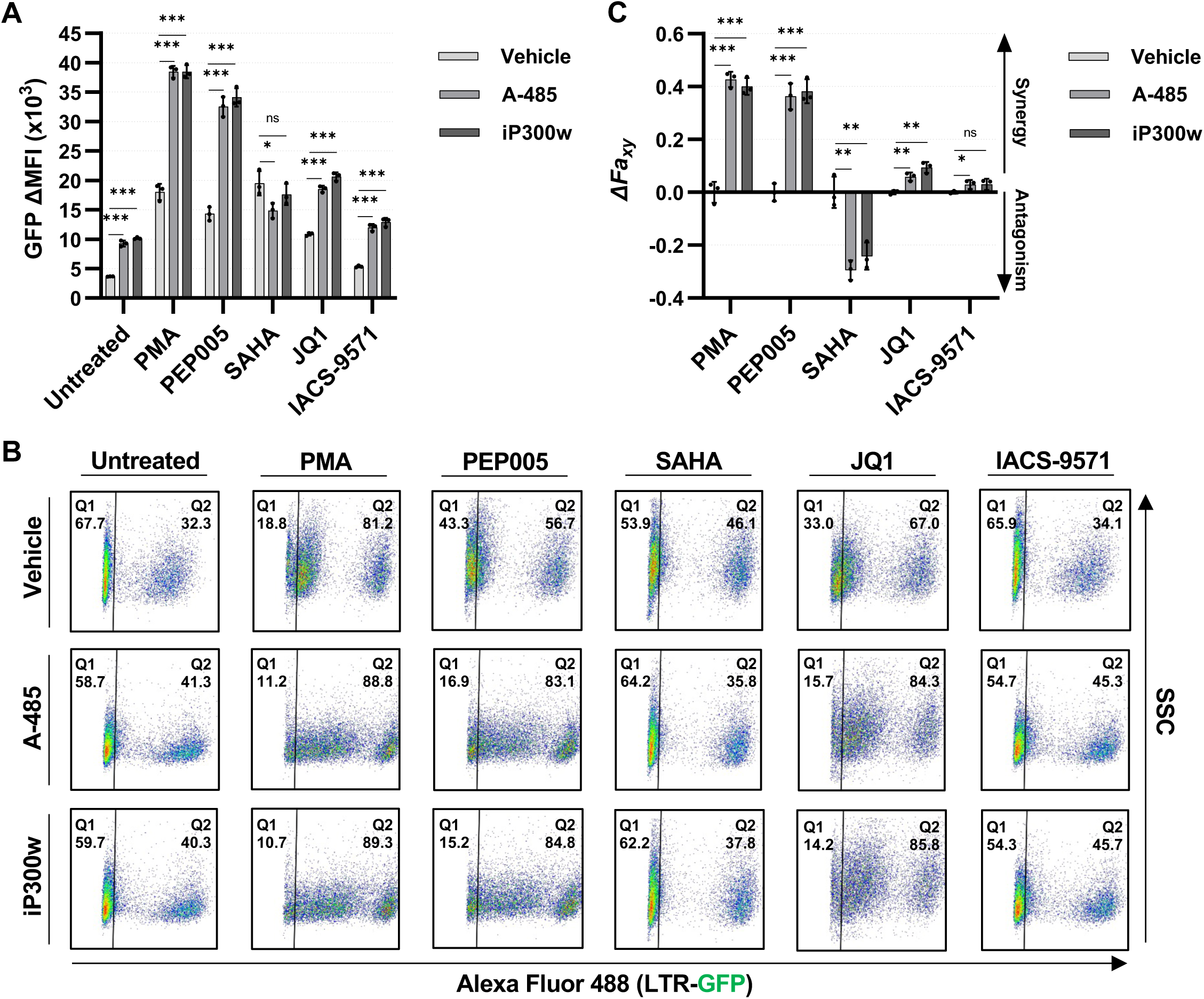
Tat dependency for CBP/p300 regulation of HIV-1. **A:** JLatA72 cells possessing an integrated HIV-1 LTR-GFP reporter that does not express Tat, were incubated 1 hr with a vehicle control (DMSO), 10 μM A-485, or 10 μM iP300w prior to the addition of 4 nM PMA, 4 nM PEP005, 1 μM SAHA, 10 μM JQ1, or 15 μM IACS-9571. Following 24 hrs, LTR transcriptional activity was assessed by flow cytometry and is reported as the GFP delta (Δ) Mean Fluorescence Intensity (MFI) (*n* = 3, mean ± SD, unpaired *t*-test). **B:** Representative flow cytometry scatter plots of JLatA72 cells treated as in (A). Cells harboring transcriptionally silent provirus (GFP-) are located in Q1 while cells with LTR expression (GFP+) are in Q2. **C:** Determination of synergy between 10 μM A-485 or μM iP300w with the indicated latency reversing agent using Bliss Independence Modelling. Drug concentrations are the same as those used in (A). Data is presented as the difference between the predicted and the observed fractional HIV-1 expression response to the given drug combination. See Materials and Methods for more details (*n* = 3, mean ± SD, unpaired *t*-test).

### Effect of CBP/p300 acetyltransferase inhibition on LTR histone modification

The above results indicate that CBP/p300 inhibition reactivates latent HIV-1 while synergizing with various mechanistically distinct LRAs to produce robust expression. However, loss of CBP/p300 function slightly impairs latency reversal in response to the HDACi SAHA. Taken together, these results indicate that CBP/p300 modulates the LTR epigenetic environment which may repress HIV-1 transcription. To examine this possibility, we performed chromatin immunoprecipitation and analysis by quantitative PCR using a set of LTR targeting primers (Fig. 7*A*). First, we performed ChIP-qPCR using non-specific IgG to determine background signal (Fig. 7*B*). Acetylation of H3K27 is catalyzed by CBP/p300 (Jin et al. 2011) (Weinert et al. 2018) (Wang et al. 2022) (Raisner et al. 2018), and unsurprisingly, CBP/p300 acetyltransferase inhibition caused a drastic reduction of H3K27ac on histones associated with all regions of HIV-1 LTR (Fig. 7*C*). Acetylation of H3K27 typically causes molecular crosstalk resulting in tri-methylation of H3K4 (Crump et al. 2011) (Tang et al. 2013) (Zhao et al. 2021). As such, we examined the effect of CBP/p300 inhibition on the establishment of LTR-associated H3K4me3. In correlation with the loss of H3K27ac, CBP/p300 inhibition caused reduced association of H3K4me3 with the HIV-1 promoter (Fig. 7*D*). Recent studies have indicated that histone H4 acetylation recruits BRD4 to the LTR resulting in reduced proviral transcription, which may provide a potential mechanism for acetylation-based inhibition of HIV-1 transcription (Li et al. 2018) (Hsieh et al. 2023). However, we observed no change in association of BRD4 with the LTR in response to CBP/p300 inhibition (Fig. 7*E*). Finally, we found that treatment with the CBP/p300 inhibitors in combination with PEP005, a PKC agonist, caused substantial recruitment of RNAPII and Serine 2 phosphorylated (S2P) RNAPII to the HIV-1 promoter (Fig. 7*F*, 7*G*). Phosphorylation of the Serine 2 residue of RNAPII occurs during elongation, however, given that the enrichment of RNAPII S2P is proportional to that of RNAPII, the predominant effect of CBP/p300 inhibition on transcription from the LTR is the facilitation of RNAPII recruitment. Collectively, these results indicate that CBP/p300 inhibition causes an epigenetic transformation that uniquely primes HIV-1 provirus for transcriptional activation.

**Figure 7.**
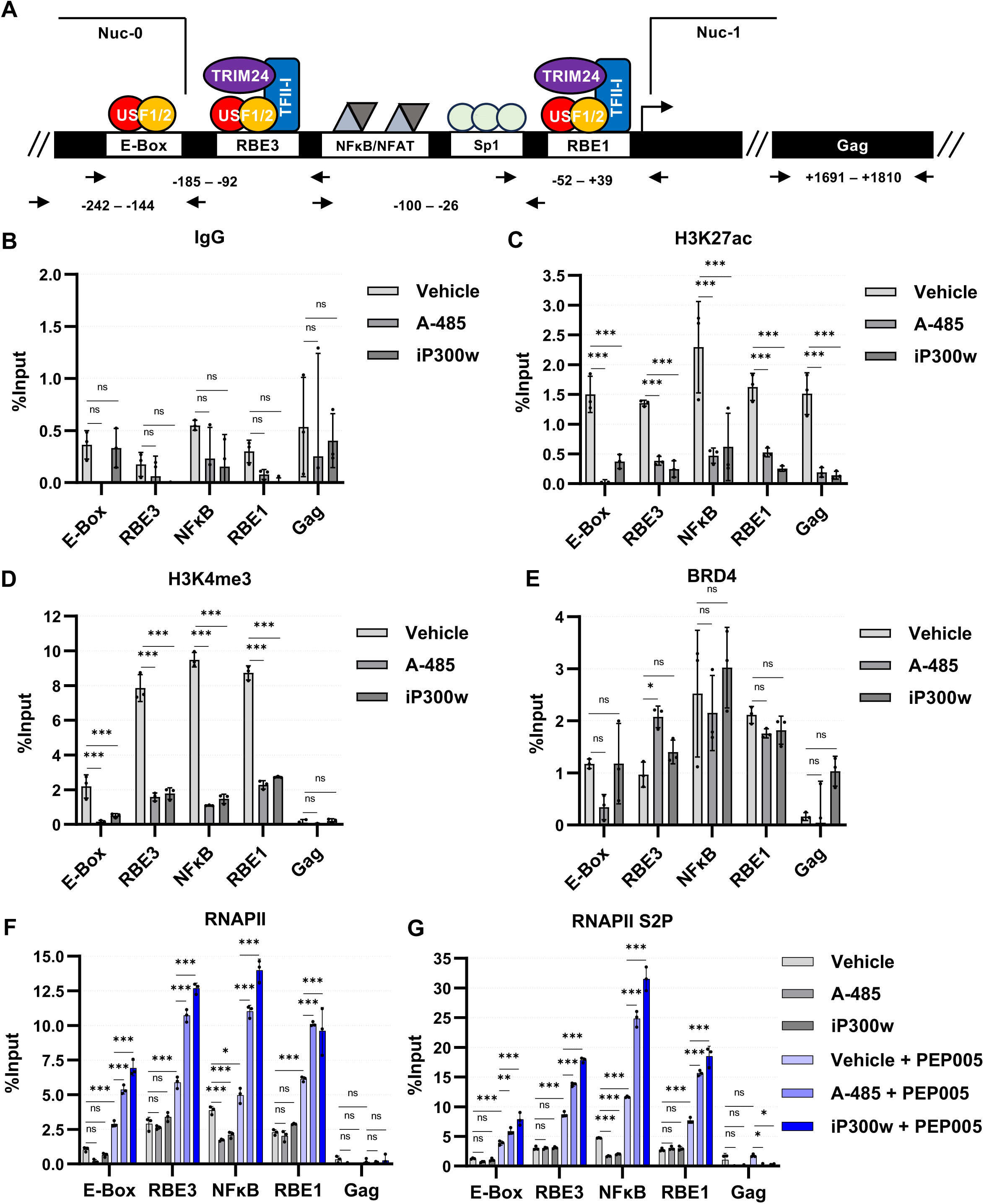
Effect of CBP/p300 inhibition on LTR epigenetic context. **A:** Schematic representation of the HIV-1 LTR and the primer pairs used for ChIP-qPCR analysis. **B:** JLat10.6 cells were incubated with a vehicle control (DMSO), 10 μM A-485, or 10 μM iP300w for 24 hrs. Subsequently, ChIP was performed using non-specific rabbit IgG and the pulled down chromatin was analyzed by qPCR (*n* = 3, mean ± SD, Two-way ANOVA with Dunnett’s multiple comparisons test). **C - E:** Following treatment of JLat10.6 cells as stated in (B), ChIP was performed using antibodies specific to H3K27ac (C), H3K4me3 (D), or BRD4 (E). Subsequently, enrichment of chromatin was determined by qPCR using oligos directed at the indicated region (*n* = 3, mean ± SD, Two-way ANOVA with Dunnett’s multiple comparisons test). **F, G:** JLat10.6 cells were incubated 24 hrs with a vehicle control (DMSO), 10 μM A-485, or 10 μM iP300w or were pre-treated 1 hr with a vehicle, 10 μM A-485, or 10 μM iP300w prior to the addition of 4 nM PEP005 followed by incubating overnight. Afterwards, ChIP was performed using antibodies specific to RNAPII (F) or S2P modified RNAPII (G), and chromatin was assessed by qPCR (*n* = 3, mean ± SD, Two-way ANOVA with Dunnett’s multiple comparisons test).

### Inhibition of PCAF/GCN5 or KAT6A/KAT6B reactivate latent HIV-1

The above results point to the unexpected conclusion that CBP/p300 has a repressive function for expression from the HIV-1 LTR. It is thought that additional acetyltransferases, including PCAF and GCN5, are important for the maintenance of productive infections (Lusic et al. 2003). To determine the role of these additional acetyltransferases for the regulation of HIV-1 transcription, we used GSK4027, a highly specific bromodomain inhibitor of PCAF/GCN5 (Humphreys et al. 2017), and WM-1119, a highly specific KAT6A/KAT6B acetyltransferase inhibitor (Baell et al. 2018) (Fig. 8*A*). Treatment of the JLat10.6 HIV-1 cell line with either inhibitor caused a dose-dependent increase in expression of GFP from the LTR, although treatment with the PCAF/GCN5 inhibitor GSK4027 caused a greater effect than the KATA/KATB inhibitor WM-1119 (Fig. 8*B*, 8*C*). Notably, reactivation in response to these inhibitors was markedly less than that caused by CBP/p300 inhibition (Fig. 1*B*, 1*C*, 3*B*). Next, we examined the effect of GSK4027 and WM-1119 in combination with various LRAs. Here, we found that combining either acetyltransferase inhibitor with PMA, PEP005, JQ1, or IACS-9571 resulted in enhanced viral expression as compared to either treatment alone (Fig. 8*D*, 8*E*). Interestingly, PCAF/GCN5 inhibition had no effect on latency reversal in response to the HDACi SAHA, while KAT6A/KAT6B hindered proviral reactivation (Fig. 8*D*, 8*E*, SAHA). Application of Bliss Independence Modeling confirmed synergistic interactions between the acetyltransferase inhibitors and PMA, PEP005, JQ1, and IACS-9571, while showing antagonism towards SAHA (Fig. 8*F*). Of note, none of the treatments caused cellular toxicity (Fig. S6). Taken together, these results indicate that acetyltransferases have more complex functions than to promote transcriptional activation of HIV-1 expression in T-cells.

**Figure 8.**
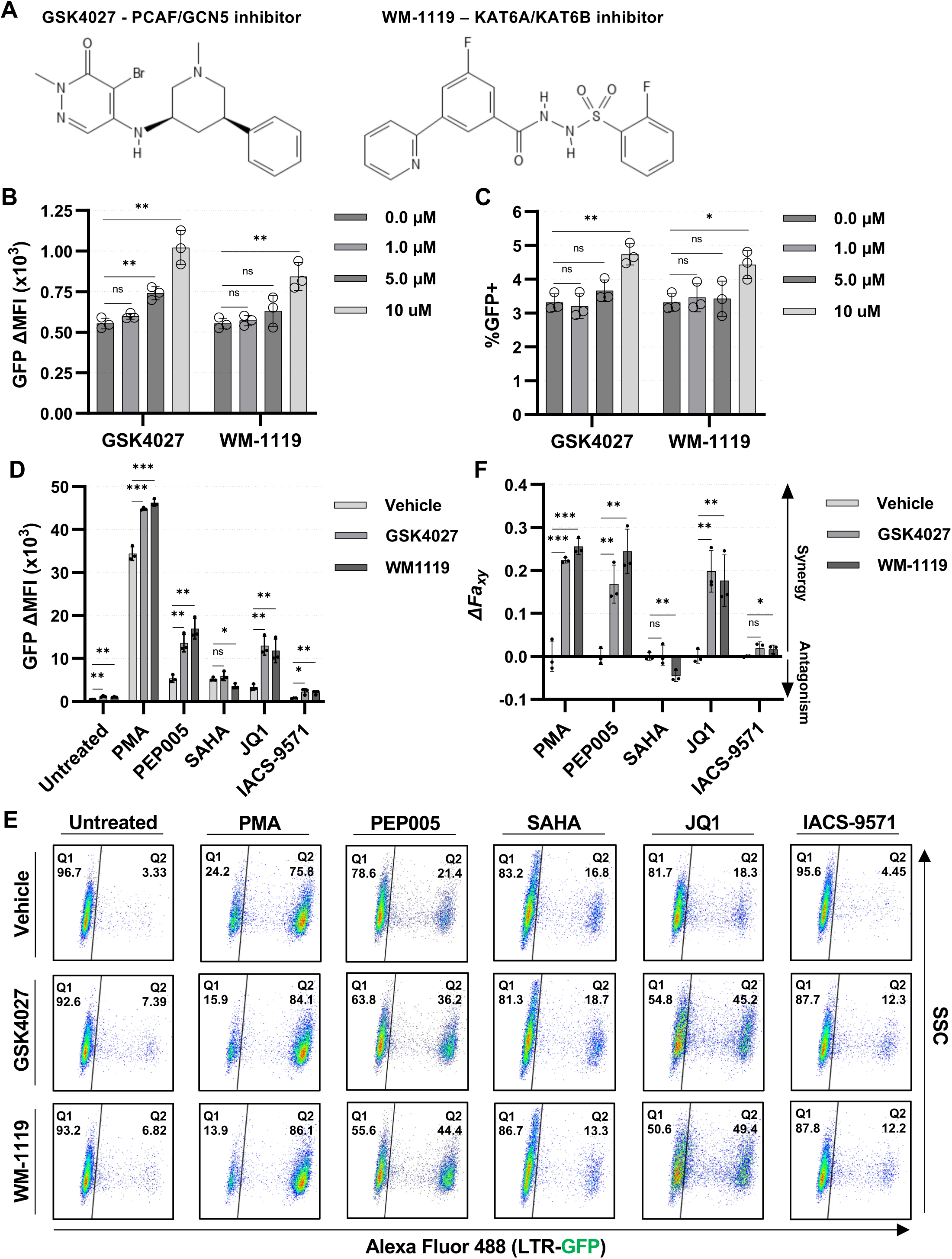
Inhibitors of various acetyltransferases induce proviral transcription. **A:** Chemical structure of GSK4027, a highly selective PCAF/GCN5 bromodomain inhibitor, and WM-1119, a highly selective KAT6A/KAT6B acetyltransferase inhibitor. **B, C:** JLat10.6 cells were incubated with the indicated concentration of GSK4027 or WM-1119. Following 24 hrs, HIV-1 transcription was assessed by flow cytometry and is stated as the delta (Δ) GFP Mean Fluorescence Intensity (MFI) (B) and the percentage of GFP positive cells (C) (*n* = 3, mean ± SD, unpaired *t*-test). **D:** JLat10.6 cells were pre-treated with a vehicle control (DMSO), 10 μM GSK4027, or 10 μM WM-1119 for 1 hr prior to the addition of the indicated latency reversing agent. After 24 hrs, HIV-1 expression was assessed by flow cytometry and is reported as the GFP delta (Δ) Mean Fluorescence Intensity (MFI). The concentration of LRAs used were 4 nM PMA, 4 nM PEP005, 1 μM SAHA, 10 μM JQ1, 15 μM IACS-9571 (*n* = 3, mean ± SD, unpaired *t*-test). **E:** Representative flow cytometry scatter plots for JLat10.6 cells treated as indicated in (D). The latent population is in Q1 (GFP-) while the transcriptionally active population is shown in Q2 (GFP+). **F:** Bliss Independence Modeling was applied to determine combinatorial drug interaction. Values greater than 0 indicate synergy while values less than 0 signify an antagonistic relationship (*n* = 3, mean ± SD, unpaired *t*-test).

## Discussion

Combination antiretroviral therapy (cART) has greatly improved the prognostic outcome of PLWH. However, cART does not represent a cure, and a cocktail of drugs must be administered for the entirety of their life for infected individuals (Chun et al. 2000) (SMART Study Group et al. 2006). Given the persistence of HIV-1, recent research efforts have examined potential strategies capable of eliminating the need for continual antiretroviral treatment such as the ‘shock and kill’ and ‘block and lock’ approaches which would involve treatment with latency reversing agents (LRAs) and latency promoting agents (LPAs), respectively (Lewis et al. 2023). Although a number of clinical trials have been performed using LRAs, none have successfully reduced the population of latently infected T-cells (Sadowski & Hashemi, 2019). Additionally, a variety of LPAs that show promising results in various HIV-1 cell line models have been identified, although viral rebound occurs at various times following removal of treatment, and no agent has proceeded to clinical trials (Mediouni et al. 2022). In general, both of these potential curative therapeutic strategy require further characterization of the mechanisms that regulate HIV-1 latency.

Stochastic HIV-1 reactivation is thought to occur in part due to fluctuations in associated histone acetylation which causes chromatin opening (Ne et al. 2018). With the goal of producing epigenetically silenced HIV-1 provirus to induce deep latency, we targeted the acetyltransferase activity of CBP/p300 using the highly specific inhibitors A-485 and iP300w. Much to our surprise, small molecule inhibition of CBP/p300 acetyltransferase function resulted in reversal of proviral latency *in vitro* (Fig. 1) and in CD4^+^ participant samples *ex vivo* (Fig. 4). To rule out the possibility that the resultant upregulation was derived from unexpected off target effects, we used the PROTAC dCBP-1 to degrade CBP and p300 via the ubiquitin/proteasome pathway. Targeted degradation of CBP/p300 caused effects that mirrored that of acetyltransferase inhibition, resulting in a dose-dependent increase in proviral transcription (Fig. 3). Furthermore, we found that CBP/p300 inhibitors produce synergistic effects with LRAs whose mechanism of action involves T-cell signaling, BRD4 bromodomain inhibition, or TRIM24 bromodomain inhibition, while antagonizing SAHA mediated reactivation (Fig. 2, 3). Consistent with the finding that CBP/p300 inhibition limits latency reversal in response to HDACi, we found that CBP/p300 activity contributes to placement of H3K27ac and H3K4me3 at the LTR (Fig. 7). Furthermore, small molecule inhibitors of the acetyltransferases PCAF/GCN5 and KAT6A/KAT6B were also shown to have latency reversal activity (Fig. 8), although to a lesser extent than CBP/p300 inhibitors.

Acetylation of histone N-terminal tail lysine residues neutralizes the ε-amino group resulting in decompaction of adjacent nucleosomes and are epigenetic marks typically associated with transcriptional activation (Shvedunova & Akhtar, 2022). A number of factors, including YY1, LSF, and NFκB p50 homodimers bind to the LTR and recruit a variety of redundant HDACs leading to hypoacetylation and suppression of HIV-1 transcription (Verdikt et al. 2021) (Peterson et al. 2023). Consistent with the role of HDACs for proviral silencing, HDACi’s have long been known to reverse latency by causing global histone acetylation. This understanding led to a focus on HDACi’s as latency reversing agents for ‘shock and kill’ clinical trials with a number of trials performed over the past two decades (Wightman et al. 2012). However, the results of these have not been encouraging as they failed to reduce the size of the latent viral reservoir despite causing a detectable increase in HIV-1 mRNA (Schou et al. 2023). The work presented here demonstrates that CBP/p300 in addition to other acetyltransferases including PCAF, GCN5, KAT6A, and KAT6B, repress HIV-1 transcription. This finding has important implications regarding the use of HDACi’s for ‘shock and kill’ strategies as although HDAC inhibition succeeds in reactivating a distinct proviral population, the balance between protein acetylation and deacetylation is likely more nuanced for the regulation of HIV-1 expression. Furthermore, while a subset of cellular genes is repressed, CBP/p300 acetylation more frequently activates cellular gene expression (Weinert et al. 2018). Given the unique suppressive effect of CBP/p300 on HIV-1, this indicates that the retrovirus has evolved to produce a stochastic expression pattern that is unique from the majority of cellular genes.

Mechanistically, we observe that CBP/p300 inhibition reduces LTR associated H3K27ac and H3K4me3 (Fig. 7*C*, Fig. 7*D*). A large number of diverse proteins bind chromatin through reader domains including bromodomains, plant homeodomains, and YEATS domains (Arrowsmith & Schapira, 2019). It is possible that CBP/p300 acetyltransferase activity creates an epigenetic context that either enables recruitment of transcriptional activators or displaces transcriptional repressors from the HIV-1 LTR. Recent studies have shown that acetylation of H4 causes recruitment of BRD4 which inhibits proviral transcription (Li et al. 2018) (Hsieh et al. 2023). However, CBP/p300 regulation of HIV-1 is independent of the H4ac/BRD4 axis as chemical inhibition did not alter association of BRD4 with the LTR (Fig. 7*E*). CBP/p300, PCAF/GCN5, and KAT6A/KAT6B are known as canonical histone acetyltransferases which predominantly localize to the nucleus. However, they display differential acetylation of histone lysine residues; acetylation of H3K18 and H3K27 by CBP/p300 is well characterized, PCAF/GCN5 are known to target H3K9, and KAT6A/KAT6B show preference for H3K9, H3K14, and H3K23 (Shvedunova & Akhtar, 2022). As inhibitors of these acetyltransferases reverses latency, further characterization of LTR epigenetic composition will be necessary to establish whether the latency reversing effect is caused by histone modification.

In addition to histones, there are hundreds of endogenously acetylated non-histone proteins (Weinert et al. 2018) (Narita et al. 2019), and consequently histone acetyltransferases (HATs) are often referred to as lysine acetyltransferases (KATs). HAT/KAT mediated protein acetylation is involved in cellular processes including gene transcription, cell cycle regulation, DNA damage repair, and autophagy (Narita et al. 2019). In particular, non-histone protein acetylation is thought to be a major regulator of transcription as acetylation is implicated in regulating the function of over 100 transcription factors, co-factors, and nuclear receptors (Narita et al. 2019). Interestingly, protein acetylation is independent of linear sequence motif suggesting that CBP/p300 produces an ‘acetyl-spray’, modifying accessible lysine residues on proximal proteins (Weinert et al. 2018). This being the case, it is possible that CBP/p300 represses HIV-1 through acetylation of non-histone proteins. In this scenario, CBP/p300 acetylation must inhibit a transcriptional activator and/or enhance function of a transcriptional repressor.

Our results demonstrate a novel effect of HAT/KAT function for regulation of HIV-1 transcription. Because HIV-1 transcription is regulated by numerous cellular factors that cause activation or repression from the LTR, and the function of most non-histone protein acetylation has not been defined, it may be difficult to identify specific lysine acetylation(s) that inhibit HIV-1 expression. Importantly, because specific protein acetylation may activate or repress HIV-1 provirus expression, successful implementation of chromatin modifying drugs for therapy will require a more detailed understanding of how protein acetylation controls function of the many factors that bind the HIV-1 LTR promoter.

## Materials and Methods

### Cell culture, virus culture, and lentiviral transduction

Jurkat E6-1, JLat10.6, JLatA72, cell lines were cultured in Roswell Park Memorial Institute 1640 medium (RPMI-1640) supplemented with 10% Fetal Bovine Serum (FBS), penicillin [100 units/ml], streptomycin [100 g/ml], and L-glutamine [2 mM]. HEK293T cells were cultured in Dulbecco’s modified Eagle’s medium (DMEM) supplemented with 10% FBS, penicillin [100 units/ml], streptomycin [100 g/ml], and L-glutamine [2 mM]. All tissue cultures were incubated in a humidified 37°C and 5% CO_2_ atmosphere.

Peripheral Blood Mononuclear Cells (PBMCs) from participants with HIV-1 on ART were isolated from whole blood by density gradient centrifugation using Lymphoprep^TM^ and SepMate^TM^ tubes (StemCell Technologies), and cryopreserved. Upon thawing, PBMCs were cultured in RPMI supplemented with 10% FBS, penicillin [100 units/ml], streptomycin [100 g/ml], L-glutamine [2 mM], and 40 U/mL IL2. Samples from participants were collected with written informed consent under a protocol jointly approved by the research ethics boards at Providence Health Care/UBC and Simon Fraser University (certificate H16-02474).

Vesicular stomatitis virus G (VSV-G) pseudotyped viral stocks were produced by co-transfecting HEK293T cells with a combination of viral molecular clone, psPAX, and VSV-G at a ratio of 8 μg: 4 μg: 2 μg. Transfections were performed with polyethylenimine (PEI) at a ratio of 6: 1 (PEI: DNA) in Gibco Opti-MEM^TM^. Lentiviral infections were performed by plating 1×10^6^ cells in 24-well plates with 500 μL RPMI containing 8 μg/mL polybrene and the amount of viral supernatant to give the desired multiplicity of infection (MOI) as indicated. Plates were spinoculated for 1.5 hrs at 1500 rpm.

Human Peripheral Blood CD4^+^ T-cells, purchased from STEMCELL Technologies (Catalog #200-0165, Lot #2107423002), were cultured in RPMI supplemented with 10% FBS, penicillin [100 units/ml], streptomycin [100 g/ml], L-glutamine [2 mM], and 40 U/mL IL2. For RGH infection, T-cells were incubated with Dynabeads^TM^ Human T-Activator CD3/CD28 beads for three days. Subsequently, the beads were removed, 8 μg/mL polybrene and RGH virus was added, and cells were spinoculated for 2 hrs at 1500 rpm and 32°C.

### Cell Viability

Cells were stained with Gibco^TM^ Trypan Blue Solution (0.4%) and viability was determined using a Bio-Rad TC20 Automated Cell Counter.

### Immunoblotting

Cells treated as indicated in the figure legends were lysed in RIPA buffer (50 mM Tris-HCl pH 7.5, 150 mM NaCl, 1% Triton X-100, 1 mM EDTA, 0.1% SDS, 0.5% sodium deoxycholate, 2.5 mM PMSF) at 4°C. Equivalent amounts of cellular extracts were mixed with 5× SDS–PAGE sample buffer, boiled for 3 min, separated using SDS–PAGE gel, and transferred onto a nitrocellulose membrane. The membrane was blocked with 2% milk (w/v) in TBS with 0.1% Tween for 1 hour at room temperature and then incubated with a primary antibody overnight at 4°C. Subsequently, the membrane was incubated for 1 hour at room temperature with a secondary HRP antibody in 2% milk (w/v) in TBS with 0.1% Tween after which the autoradiograph was developed. Antibodies used are as follows: Tubulin, Abcam ab7291, 1:13333; p300, Cell Signaling Technology #54062, 1:1000; CBP, Abcam ab253202, 1:2000; Goat Anti-Rabbit-HRP, Abcam ab6721, 1:1100000; Goat Anti-Mouse-HRP, Pierce 1858413, 1:20000.

### RT-PCR

RNA was extracted from cells using the RNeasy Kit (Qiagen) and analyzed with the Quant Studio 3 Real-Time PCR system (Applied Biosystems) using *Power* SYBR® Green RNA-to-CT™ 1-Step Kit (Thermo Fisher) as per the manufacturer’s instructions. RT-PCR data was normalized to GAPDH expression using the ΔΔCt method as previously described ( Livak & Schmittgen, 2001). Cycling parameters were as follows: 48 °C, 30 min, 1x; 95 °C, 10 min, 1x; (95 °C, 15 sec, 60 °C, 1 min), 50x. Primers were as follows: HIV-1 mRNA (multiply spliced Tat-Rev transcripts), Fwd 5’ CTTAGGCATCTCCTATGGCAGGA, Rev 5’ GGATCTGTCTCTGTCTCTCTCTCCACC; *GAPDH*, Fwd 5’ TGCACCACCAACTGCTTAGC, Rev 5’ GGCATGGACTGTGGTCATGAG; *IL2*, Fwd 5’ AACTCACCAGGATGCTCACA, Rev 5’ GCACTTCCTCCAGAGGTTTGA; *CD69*, Fwd 5’ TCTTTGCATCCGGAGAGTGGA, Rev 5’ ATTACAGCACACAGGACAGGA; *IL8*, Fwd 5’ ACTGAGAGTGATTGAGAGTGGAC, Rev 5’ AACCCTCTGCACCCAGTTTTC; *CSF2*, Fwd 5’ ACCTGCCTACAGACCCGCCT, Rev 5’ GAAGTTTCCGGGGTTGGAGGGC; *CCL22*, Fwd 5’ CGCGTGGTGAAACACTTCTAC, Rev 5’ GCCACGGTCATCAGAGTAGG; *XCL1*, Fwd 5’ CTCCTTGGCATCTGCTCTCT, Rev 5’ GCTCACACAGGTCCTCTTATC.

### Flow cytometry

Cells were treated as indicated in the figure legends and suspended in PBS. Flow cytometry was performed using a BD Biosciences LSRII-561 system with threshold forward scatter (FSC) and side scatter (SSC) parameters set so that a homogenous population of live cells was assessed. FlowJo software (TreeStar) was used to analyze data and determine the indicated Mean Fluorescence Intensity (MFI). GFP delta (Δ) Mean Fluorescence Intensity (MFI) specifies that the MFI of the Jurkat E6-1 negative control sample has been subtracted.

### Quantitative analysis of drug interactions

As previously described (Laird et al. 2015) (Liu et al. 2018), Bliss independence modeling was applied to assess the interaction of the indicated acetyltransferase inhibitor with the indicated LRA in relationship to HIV-1 expression. The Bliss independence model is defined by the equation *Fa*_xy, P_ = *Fa_x_* + *Fa_y_* – (*Fa_x_*)(*Fa_y_*), whereby *Fa*_xy, P_ is the predicted fraction affected by a combination of drug *x* and drug *y* that is derived from the experimentally observed fraction affected by drug *x* (*Fa_x_*) and drug *y* (*Fa_y_*) individually. Comparison of the predicted combinatorial affect (*Fa*_xy, P_) with the experimentally observed impact (*Fa_xy_*_,*O*_) is then performed: Δ*Fa_xy_* = *Fa_xy_*_,*O*_ − *Fa_xy_*_,*P*_. If Δ*Fa_xy_* is greater than 0, the combination of drugs *x* and *y* exceed that of the predicted affect indicating that the drugs possess synergistic interaction. If Δ*Fa_xy_* = 0, the drug combination follows the Bliss model for independent action. If Δ*Fa_xy_* is less than 0, the drug interaction is antagonistic as the observed effect of the drug combination is less than predicted combined affect. In this analysis, the fraction affected was calculated as follows: *Fa_x_* = (GFP ΔMFI of drug *x* – GFP ΔMFI untreated) / (maximum GFP ΔMFI obtained by any given treatment within the experiment – GFP ΔMFI untreated).

### ChIP-qPCR

JLat10.6 cells were fixed with 1% formaldehyde (Sigma-Aldrich) for 10 min at room temperature (1.5×10^7^ cells per ChIP). Crosslinking was quenched by the addition of 125 mM glycine for 5 min, after which cells were washed with PBS at 4°C. Cells were lysed in NP-40 Lysis Buffer (0.5% NP-40, 10 mM Tris-HCl pH = 7.8, 3 mM MgCl_2_, 1x PIC, 2.5 mM PMSF) for 15 min on ice. Following sedimentation, nuclei were resuspended in Sonication Buffer (10 mM Tris-HCl pH = 7.8, 10 mM EDTA, 0.5% SDS, 1x PIC, 2.5 mM PMSF) and sonicated using a Covaris S220 Focused-ultrasonicator to produce sheared DNA that predominately ranged from 200 – 2000 bp using the following settings: Treatment 1 [ Peak Power: 200, Duty Factor: 10, Cycles/Burst 200, Duration 30 sec]; Treatment 2 [ Peak Power: 2.5, Duty Factor: 0.1, Cycles/Burst 50, Duration: 30 sec]; Repeat 6x. The soluble chromatin fraction was collected and snap frozen in liquid nitrogen. Chromatin concentrations were normalized among samples and pre-cleared with Protein G agarose beads (Millipore, 100 μL). The chromatin samples were split in two and diluted with IP buffer (10 mM Tris-HCl pH = 8.0, 1.0% Triton X-100, 0.1% deoxycholate, 0.1% SDS, 90 mM NaCl, 2 mM EDTA, 1x PIC); samples were immunoprecipitated with the indicated specific antibody or run as a no antibody mock IP. Antibodies used for ChIP are as follows: Rabbit IgG, Abcam ab1722730, 10 μg; H3K27ac, Abcam ab4729, 4 μg; H3K4me3, Abcam ab213224, 3 μg; BRD4, Abcam ab314432, 10 μg; RNAPII, Abcam ab26721, 4 μg; RNAPII S2P, Abcam ab238146, 5 μg.

The chromatin/ antibody mixtures were incubated 1 hr at 4°C with rotation. Pre-washed Protein G agarose beads (40 μL) were then added, and the samples were incubated overnight at 4°C with rotation. Bead/antibody complexes were washed 3x in Low Salt Wash Buffer (20 mM Tris-HCl pH = 8.0, 0.1% SDS, 1.0% Triton X-100, 2 mM EDTA, 150 mM NaCl, 1x PIC) and 1x with High Salt Wash Buffer (same as Low Salt but with 500 mM NaCl). Elution and crosslink reversal was performed by incubating 4 hrs at 65°C in elution buffer (100 mM NaHCO_3_, 1% SDS) supplemented with RNase A. DNA was purified using the QIAQuick PCR purification kit (QIAGEN) and ChIP DNA was analyzed using the Quant Studio 3 Real-Time PCR system (Applied Biosystems). The percent input value of the sample paired no antibody mock immunoprecipitation has been subtracted from the percent input obtained from immunoprecipitation with IgG or the indicated specific antibody of the corresponding sample. Oligos used for ChIP-qPCR are as follows: E-Box, Fwd 5’ GTGAGCCTGCATGGAATGGA, Rev 5’ CGGATGCAGCTCTCGGG; RBE3, Fwd 5’ AGCCGCCTAGCATTTCATC, Rev 5’ CAGCGGAAAGTCCCTTGTAG; NFκB, Fwd 5’ TTTCCGCTGGGGACTTTC, Rev 5’ CCAGTACAGGCAAAAAGCAG; RBE1, Fwd 5’ AGTGGCGAGCCCTCAGAT, Rev 5’ AGAGCTCCCAGGCTCAAATC; Gag, Fwd 5’ AGCAGCCATGCAAATGTTA, Rev 5’ AGAGAACCAAGGGGAAGTGA.

### RNA-seq analysis

RNA-seq as analyzed by DESeq2 to identify genes differentially expressed upon T-cell activation was obtained from NCBI GEO accession GSE227850 (Horvath & Sadowski, 2023). Volcano plots were generated using the Galaxy web platform (Afgan et al. 2018) while heatmaps depicting gene expression were created in RStudio.

### Statistics and reproducibility

All replicates are presented as mean values with ± standard deviation shown by error bars. P-values were determined using GraphPad Prism 10.2.1 with the number of replicates and the statistical method used noted in the figure legends. Statistical significance is indicated at \**P* < 0.05, \*\**P* < 0.01, or \*\*\**P* < 0.001, with n.s. denoting non-significant *P* ≥ 0.05.

## Data availability

All data supporting the findings of this study are available within the article or from the corresponding author upon reasonable request (I. Sadowski).

## Acknowledgments

We thank LeAnn Howe for helpful comments. We thank Zabrina Brumme and the BC Centre for Excellence in HIV/AIDS for providing PBMCs from study participants. We thank the laboratory staff at the BC Centre for Excellence in HIV/AIDS for processing PBMCs from study participants. This research was supported by program project grant F16-01210, from the Canadian Institutes of Health Research (CIHR), and F19-05392 Discovery Grant from the Natural Sciences and Engineering Research Council of Canada (NSERC) (to I.S.).

## Author Contributions

Horvath R. M. and Sadowski I. conceived the experimental design. Horvath R. M. performed all experiments. Horvath R. M. and Sadowski I. wrote the manuscript.

## Legends to Supplementary Figures

**Figure S1.**
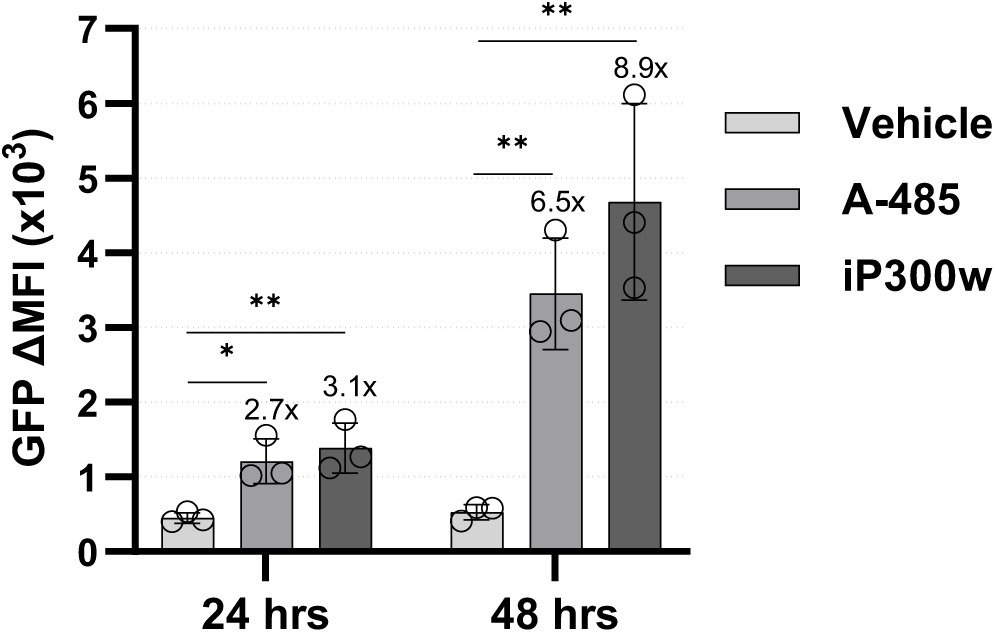
CBP/p300 inhibition causes greater proviral reactivation over time. JLat10.6 cells were incubated with a vehicle control (DMSO), 10 μM A-485, or 10 μM iP300w. Following incubation for 24 hrs or 48 hrs, proviral expression was determined by flow cytometry and is indicated as the GFP delta (Δ) Mean Fluorescence Intensity (MFI) (*n* = 3, mean ± SD, unpaired *t*-test).

**Figure S2.**
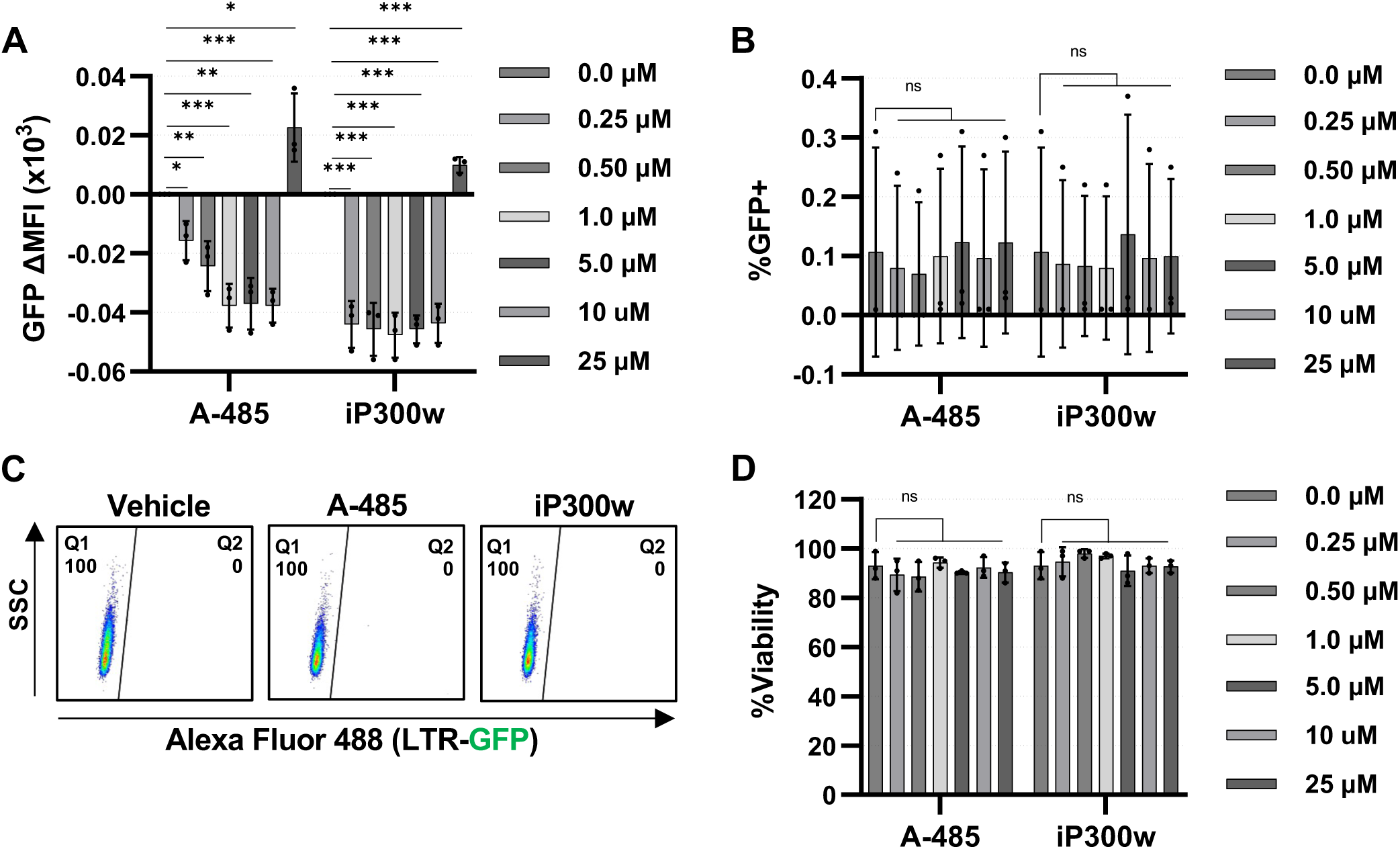
A-485 and iP300w do not cause autofluorescence. **A, B:** Jurkat E6-1 cells were incubated with the indicated concentration of A-485 or iP300w. Subsequent to 24 hrs, flow cytometry was performed to assess GFP as measured by delta (Δ) Mean Fluorescence Intensity (MFI) (A) and the percentage of GFP positive cells (B) (*n* = 3, mean ± SD, unpaired *t*-test). **C:** Representative flow cytometry scatter plots of Jurkat E6-1 cells treated as in (A, B). **D:** Viability was determined for Jurkat E6-1 cells treated as in (A, B) (*n* = 3, mean ± SD, unpaired *t*-test).

**Figure S3.**
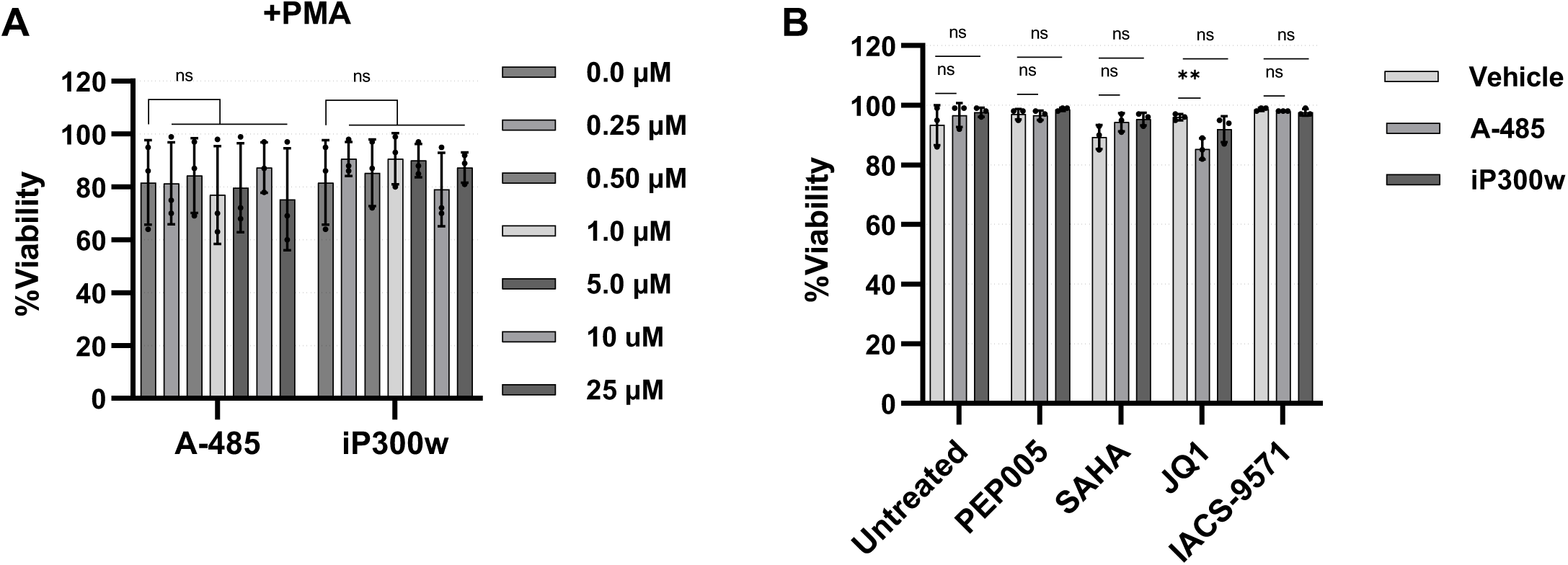
Cellular viability following treatment with CBP/p300 inhibitors in combination with LRAs. **A:** Following a 1 hr pre-treatment with the indicated concentration of A-485 or iP300w, 4 nM PMA was added to JLat10.6 cells. Cellular viability was determined after 24 hr incubation (*n* = 3, mean ± SD, unpaired *t*-test). **B:** JLat10.6 cells were pre-treated with DMSO (Vehicle), 10 μM A-485, or 10 μM iP300w for 1 hr after which the indicated LRA was added. The concentration of LRA used was 4 nM PEP005, 1 μM SAHA, 10 μM JQ1, and 15 μM IACS-9571. Viability was subsequently determined (*n* = 3, mean ± SD, unpaired *t*-test).

**Figure S4.**
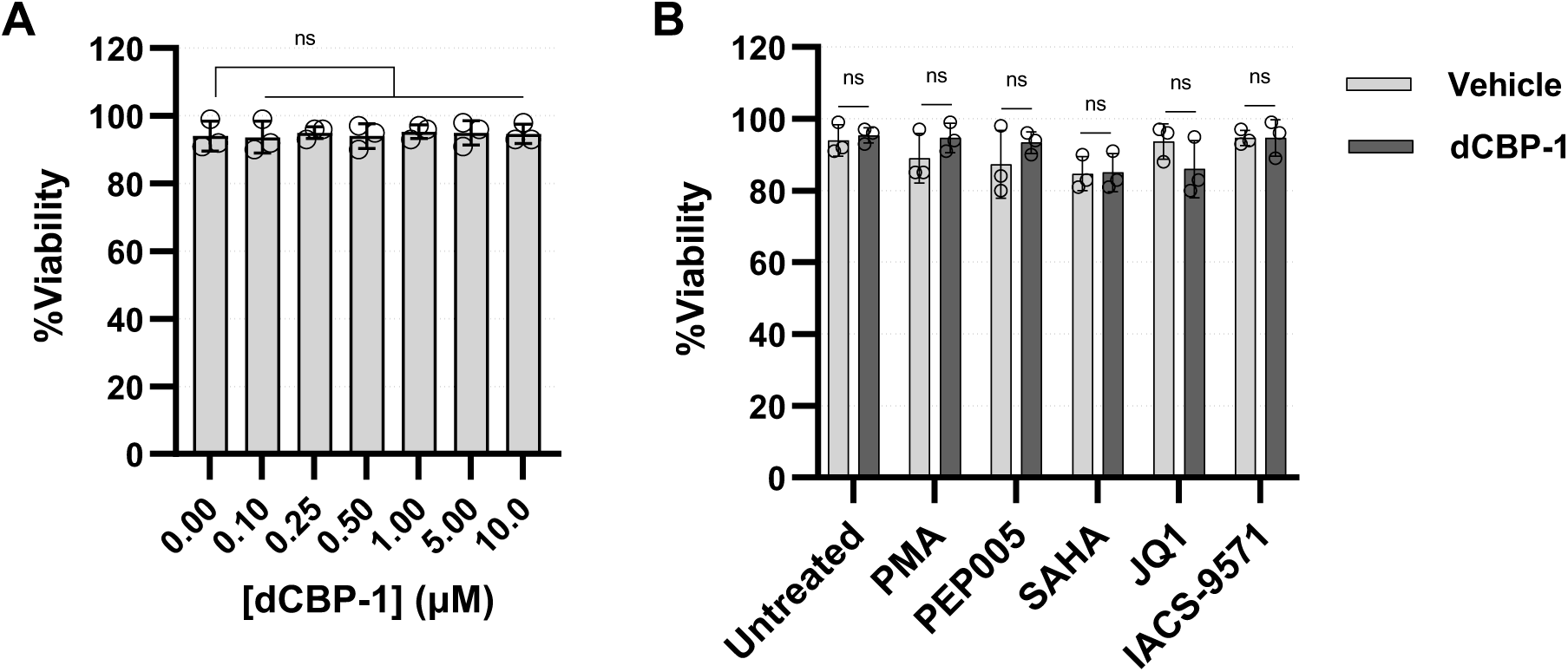
Effect of dCBP-1 on cellular viability. **A:** JLat10.6 cells were incubated with the indicated concentration of dCBP-1 for 24 hrs after which cellular viability was assessed (*n* = 3, mean ± SD, unpaired *t*-test). **B:** Following a 1 hr pre-treatment with 1 μM dCBP-1, the indicated LRA was added, and cellular viability was determined after 24 hrs. The concentration of LRA used was 4 nM PMA, 4 nM PEP005, 1 μM SAHA, 10 μM JQ1, and 15 μM IACS-9571 (*n* = 3, mean ± SD, unpaired *t*-test).

**Figure S5.**
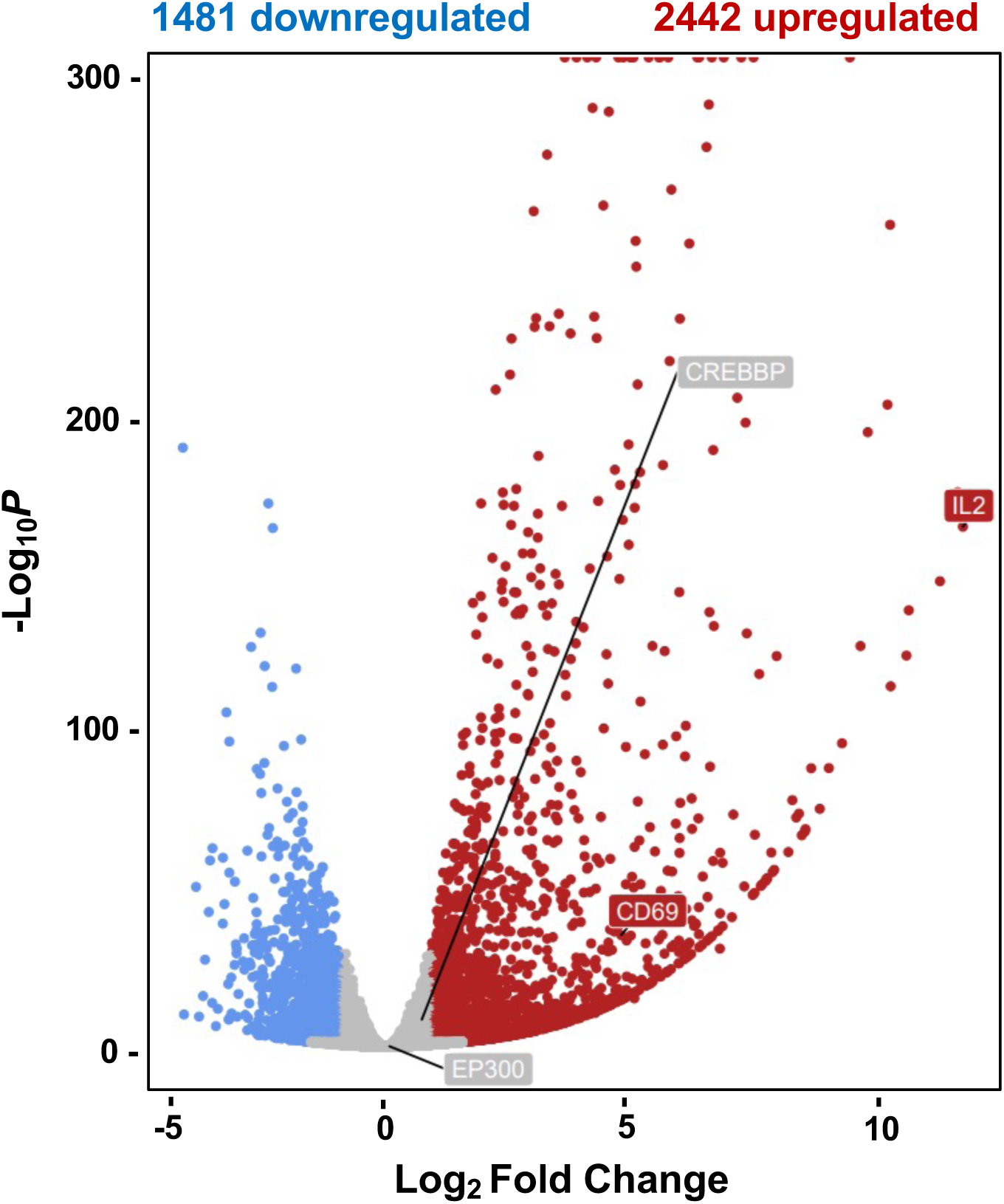
RNA-seq of activated T-cells. Volcano plot depiction of genes that are differentially expressed following the treatment of Jurkat T-cells with PMA/ionomycin as analyzed by DESeq2 (Horvath et al. 2023).

**Figure S6.**
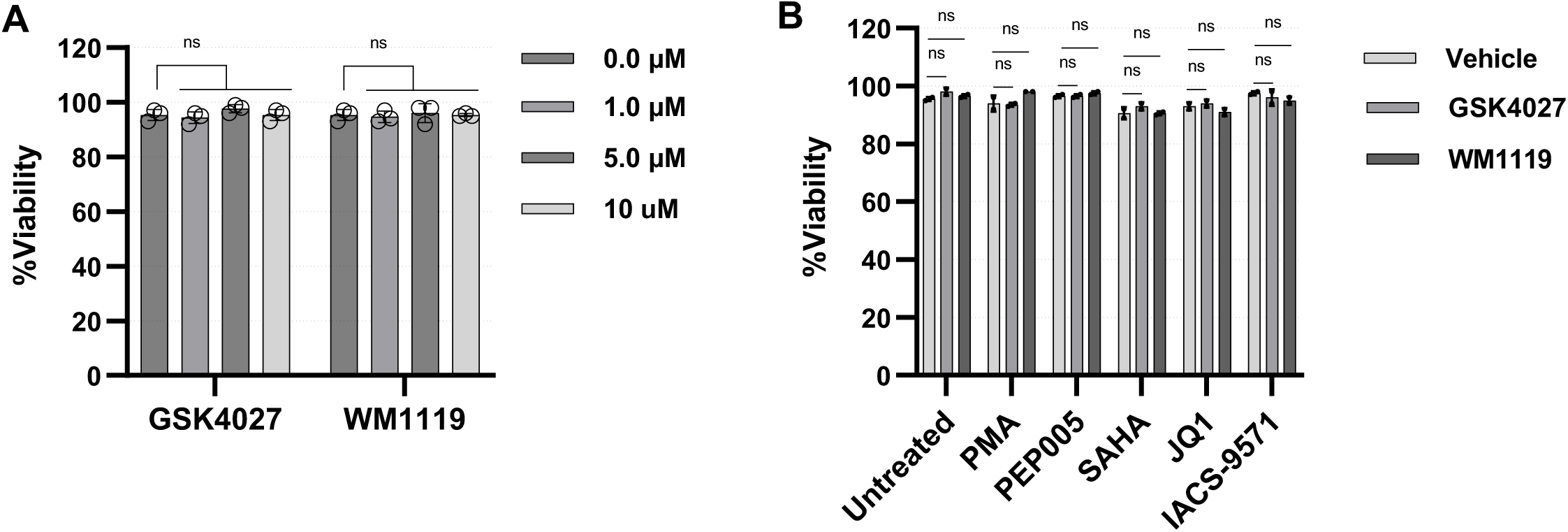
Effect of GSK4027 or WM-1119 on cellular viability. **A:** JLat10.6 cells were incubated in the presence of the indicated concentration of GSK4027 or WM-1119 for 24 hrs after which cellular viability was determined (*n* = 3, mean ± SD, unpaired *t*-test). **B:** Following a 1 hr pre-treatment with a Vehicle control (DMSO), 10 μM GSK4027, or 10 μM WM-1119, the indicated LRA was added and after 24 hrs, cellular viability determined. The concentration of LRA used was 4 nM PMA, 4 nM PEP005, 1 μM SAHA, 10 μM JQ1, and 15 μM IACS-9571 (*n* = 3, mean ± SD, unpaired *t*-test).

**Figure S7.**
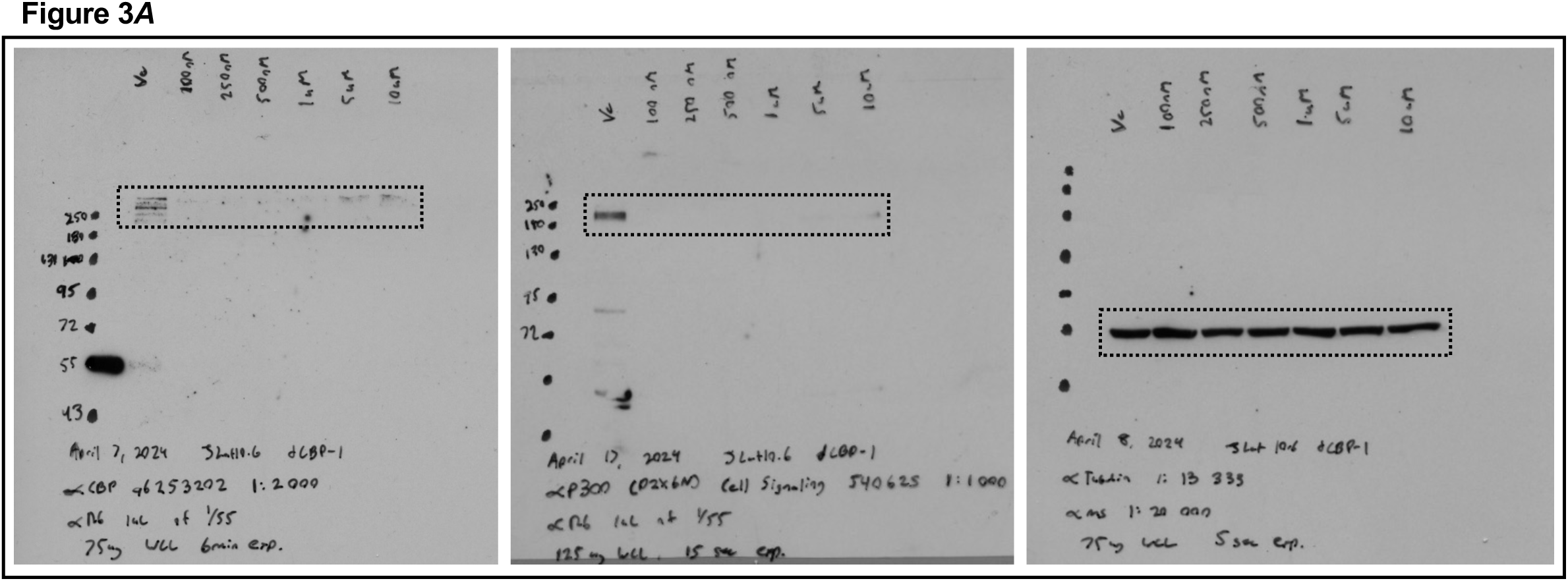
Full-length western blots.

